# Active cargo positioning in antiparallel transport networks

**DOI:** 10.1101/512863

**Authors:** Mathieu Richard, Carles Blanch-Mercader, Hajer Ennomani, Wenxiang Cao, Enrique M. De La Cruz, Jean-François Joanny, Frank Jülicher, Laurent Blanchoin, Pascal Martin

## Abstract

Cytoskeletal filaments assemble into dense parallel, antiparallel or disordered networks, providing a complex environment for active cargo transport and positioning by molecular motors. The interplay between the network architecture and intrinsic motor properties clearly affects transport properties but remains poorly understood. Here, by using surface micro-patterns of actin polymerization, we investigate stochastic transport properties of colloidal beads in antiparallel networks of overlapping actin filaments. We found that 200-nm beads coated with myosin-Va motors displayed directed movements towards positions where the net polarity of the actin network vanished, accumulating there. The bead distribution was dictated by the spatial profiles of local bead velocity and diffusion coefficient, indicating that a diffusion-drift process was at work. Remarkably, beads coated with heavy mero-myosin-II motors showed a similar behavior. However, although velocity gradients were steeper with myosin II, the much larger bead diffusion observed with this motor resulted in less precise positioning. Our observations are well described by a three-state model, in which active beads locally sense the net polarity of the network by frequently detaching from and reattaching to the filaments. A stochastic sequence of processive runs and diffusive searches results in a biased random walk. The precision of bead positioning is set by the gradient of net actin polarity in the network and by the run length of the cargo in an attached state. Our results unveiled physical rules for cargo transport and positioning in networks of mixed polarity.

**Significance statement:** Cellular functions rely on small groups of molecular motors to transport their cargoes throughout the cell along polar filaments of the cytoskeleton. Cytoskeletal filaments self-assemble into dense networks comprising intersections and filaments of mixed polarity, challenging directed motor-based transport. Using micro-patterns of actin polymerization in-vitro, we investigated stochastic transport of colloidal beads in antiparallel networks of overlapping actin filaments. We found that beads coated with myosin motors sensed the net polarity of the actin network, resulting in active bead positioning to regions of neutral polarity with a precision depending on the motor type. A theoretical description of our experimental results provides the key physical rules for cargo transport and positioning in filament networks of mixed polarity.

## Introduction

Controlling transport at microscopic scales poses a fundamental challenge in biology, engineering and physics. In the cell, molecular motors use cytoskeletal filaments as polar tracks for directed transport (1). However, the filaments are organized in dense networks with structural heterogeneities (2), which impart stringent physical constraints according to two basic mechanisms (3). First, disorder in filament orientations, as well as branching, results in intersections that can affect the directionality of cargo transport (4). Second, networks of mixed polarity foster bidirectional movements either because a given cargo can interact with multiple filaments simultaneously, resulting in a tug-of-war (5), or stochastically detach and reattach to a different filament (6, 7). Conversely, the outcome of motor-driven transport in a given cytoskeletal architecture also depends on the single-molecule properties of the motors (8–10) and on their collective organization (8, 11, 12).

Two types of theoretical approaches have been proposed to describe the collective behavior of motor assemblies attached to the same cargo. Microscopic theories aim at determining the collective behavior from single motor properties (13–16). Alternatively, coarse grained theories focus on global properties of cargo dynamics resulting from stochastic binding to and unbinding from cytoskeletal filaments (17). The latter approach introduces only a few effective parameters, such as binding and unbinding rates as well as transport velocity of the cargo, and does not rely on detailed knowledge on single motors. Thanks to its relative simplicity, the coarse-grained approach is well adapted for the study of cargo transport in complex environments (6, 7).

In this work, we developed an in-vitro assay to study the interplay between the cytoskeletal organization and motor properties for cargo transport. We monitored transport of beads that were actively propelled by either processive myosin-Va or non-processive heavy-mero myosin-II motors in antiparallel networks of actin filaments with exponential density profiles. We found that the myosin-coated beads were actively positioned to regions where the net polarity of the actin network vanished and with a higher precision than the characteristic length associated with polarity gradients of the network. A theoretical description of the transport process at the coarse-grained level was able to clarify the physical rules underlying active cargo trapping in antiparallel networks depending on motor properties and filament-polarity gradients.

## Results

### Antiparallel actin networks of controlled architecture

We designed surface micro-patterns of the nucleation-promoting factor pWA to control the geometry and collective organization of growing actin filaments (18, 19). Our patterns were composed of parallel nucleation lines spaced by a distance *L* ≈ 40 μm. Actin filaments extended perpendicularly to the nucleation lines by monomer incorporation to their barbed ends; the pointed ends of the filaments were localized on the pattern, whereas the barbed ends moved away from the nucleation zones. After ~10 min of polymerization, filaments growing in opposite directions from adjacent lines overlapped, yielding an anti-parallel network (Fig. 1). Inclusion of methylcellulose as a depleting agent ensured that most of the network remained within ~200 nm of the glass surface (20), thus forming an actin sheet.

**Figure 1:**
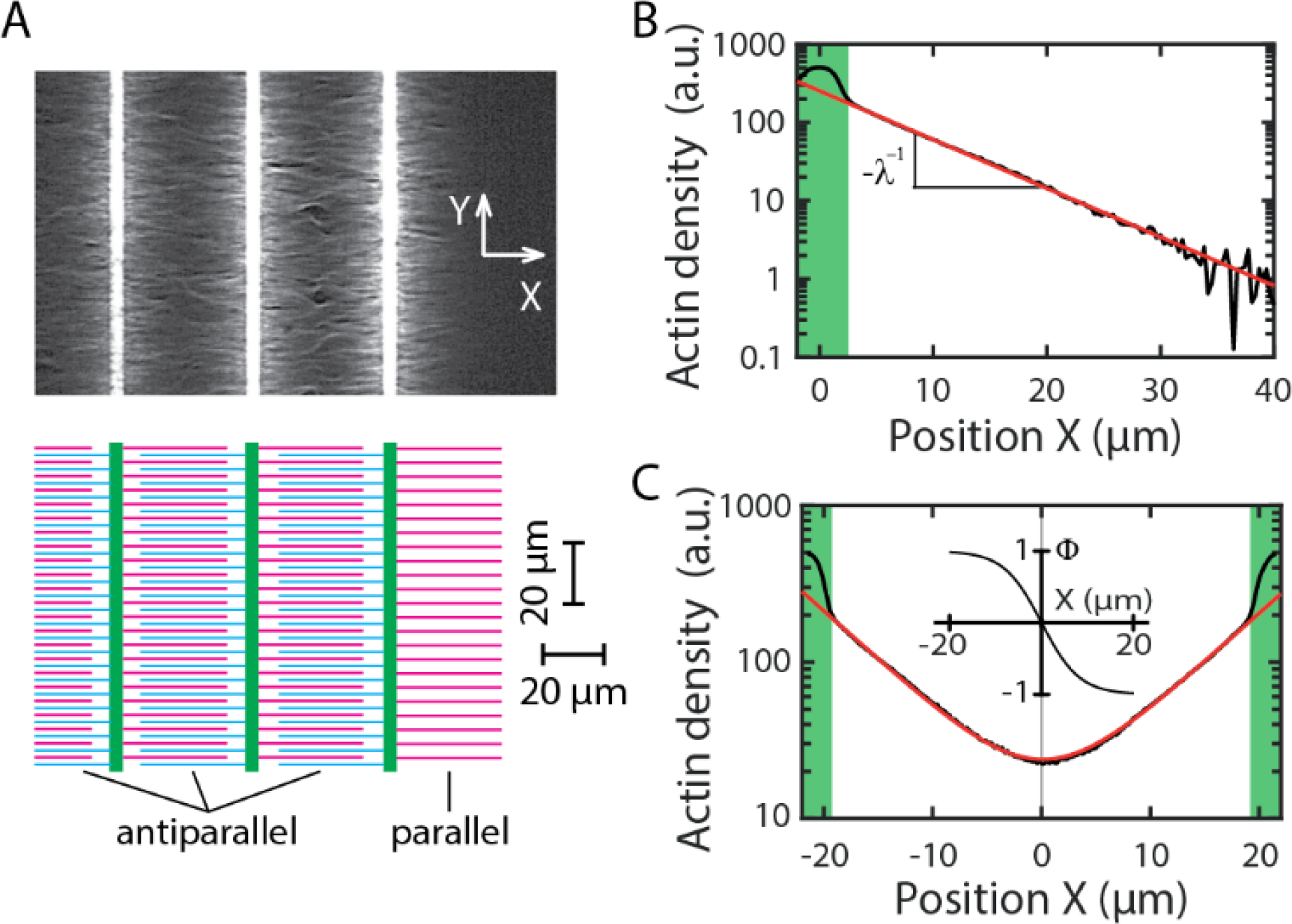
Bi-dimensional network of antiparallel actin filaments in vitro. **A:** Fluorescence imaging (top) and schematic representation (bottom) of the actin architecture. Actin filaments grew from a surface pattern of parallel lines (green) coated with a nucleation-promoting-factor of actin polymerization. The filaments are oriented along axis X, thus perpendicular to the nucleation lines (axis Y), with their barbed ends located away from the lines. An antiparallel network of overlapping actin filaments was established between neighboring nucleation lines. The line at the right end of the pattern has no neighbor on its right-hand side and thus provides a parallel network of filaments. **B:** Actin density profile for the parallel network. This semilog plot shows that the profile is well described by an exponential function (red line). The inverse absolute slope of the curve corresponds to the mean length, here *λ* = 7 μm, of the actin filaments in the network. **C:** Actin density profile for the antiparallel network. The profile is well described (red line) by the sum of two mirror-symmetric exponential with the same characteristic length as that obtained in **B**. In **B** and **C**, the nucleation lines are marked by the green shaded areas. Inset: The net polarity *ϕ*(*X*) = tanh(*X*/**λ**) of the antiparallel network is plotted as a function of position *X*, where **λ** represents the mean-actin length measured in **A**; the net polarity is null at the center of the network, where *X* = 0. Nucleation lines are spaced by 40 μm and have a width of 3 μm. The actin density profiles were measured after 30 min of polymerization.

We took advantage of the nucleation line at one edge of the pattern to characterize the architecture of a parallel actin network, without interference from actin filaments coming in opposite direction from a neighboring nucleation line. In this case, the actin density profile along an axis *X* perpendicular to the nucleation line followed an exponential decay *ρ*(*X*) ∝ exp(− *X*/*λ*) (Fig. 1B). There, the decay length *λ* = 8.2±1.1 μm (n = 29) corresponds to the mean length of actin filaments in the network.

In between two parallel nucleation lines, the actin density profile of the anti-parallel network displayed mirror symmetry about the center, where the density was minimal. In the following, the center was set at *X* = 0. Corresponding to this symmetry, the profile *ρ*(*X*) = *ρ*^+^(*X*) + *ρ*^−^(*X*) was described by the sum of two mirror-symmetric exponentials *ρ*^±^(*X*) = (*ρ*_0_/2) exp(∓*X*/*λ*); equivalently *ρ*(*X*) = *ρ*_0_ cosh(*X*/*λ*) with *ρ*_0_ the total actin density at the center (Fig. 1C). The mean filament length *λ*λ was indistinguishable from that measured with a single line at the edge of the pattern. We define the net polarity of the actin network as Φ(*X*) = (*ρ*^+^(*X*) − *ρ*^−^(*X*))/(*ρ*^+^(*X*) + *ρ*^−^(*X*)) = tanh(*X*/*λ*) (Fig. 1C, inset). At the center of the actin density profile, the net polarity vanished (Φ(0) = 0) and the net-polarity gradient was given by the inverse of the mean actin length (Φ′(0) = 1/*λ*). Thus, the shorter the average length of the filaments, the steeper the gradient of net polarity. Because the pattern spacing was significantly larger than the mean filament length (*L* ≫ *λ*), all filaments had nearly the same polarity near each of the nucleation lines (Φ(±*L*/2) = ∓1).

### Active centering of myosin-coated beads

We tracked the trajectories of 200-nm beads that were coated with either myosin Va or heavy-mero myosin II (hereafter called ‘myosin V’ or ‘myosin II’ for simplicity) after they landed on the actin network by sedimentation from the bulk. Both types of myosin-coated beads showed directed trajectories towards the midline of the network, hereafter called the center (Fig. 2A-B and E-F). Remarkably, although there were enough long filaments in the network to guide myosin-based transport all the way to the other side of the pattern (SI Appendix, Fig. S1), the beads appeared unable to move a significant distance past the center, where the net polarity of the actin network vanished, accumulating there (Fig. 2C and G). As a result, the steady-state distribution of bead positions was peaked, in contrast to the flat distribution of initial bead positions corresponding to the first detection of the beads in the network (Fig. 2D and H). We considered that beads had reached steady state when their trajectories lasted at least 200 s (Fig. 2B and F; see Methods). On average, active cargo transport by myosin V or myosin II positioned the beads precisely at the center of the antiparallel actin network.

**Figure 2:**
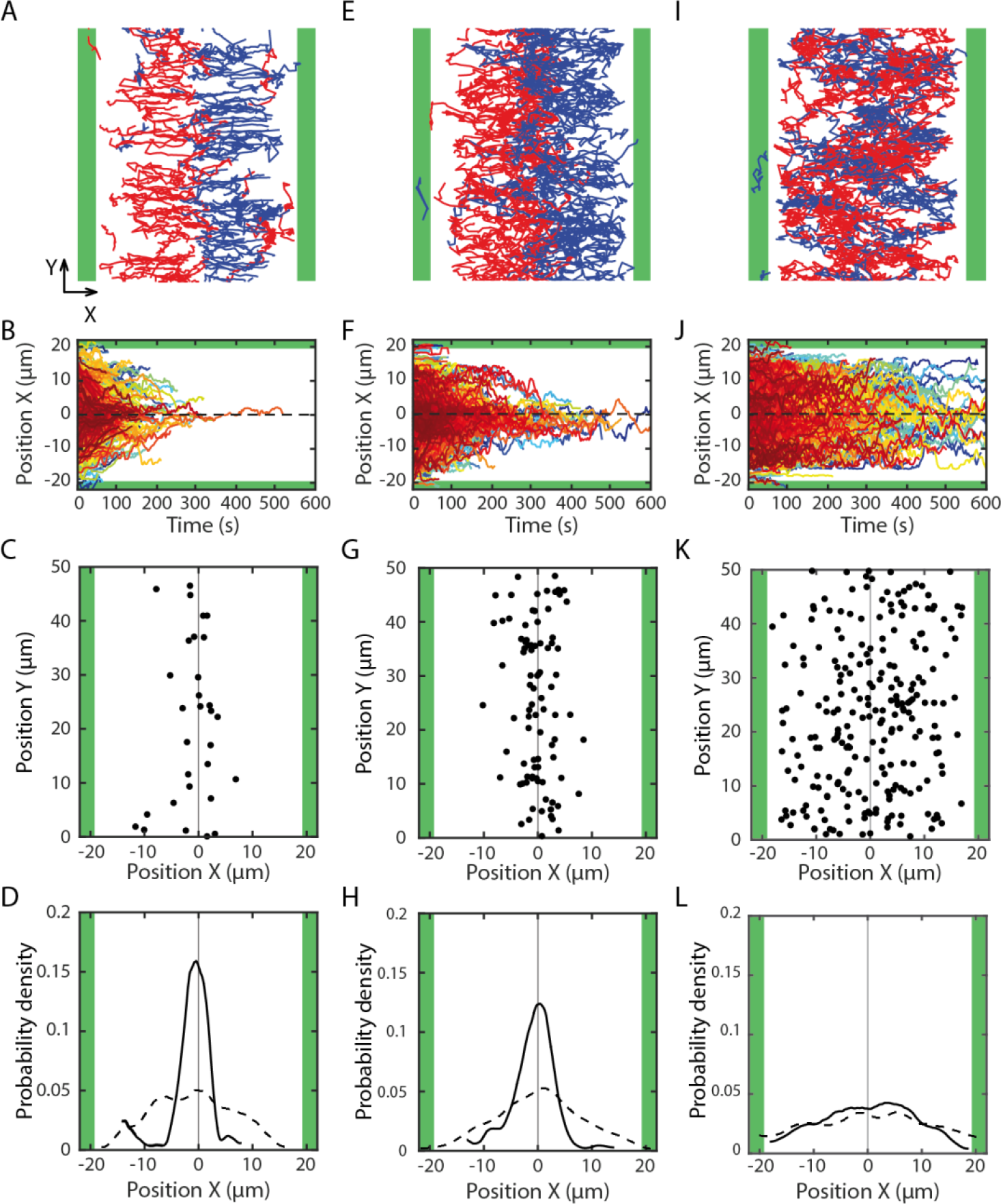
Bead transport and steady-state distribution in antiparallel actin networks. Beads were coated with molecular motors (myosin Va in **A-D**; myosin II HMM in **E-H**) or passivated with BSA (**I-L**). **A, E** and **I**: Bead trajectories within the plane of the actin network. Red (blue): trajectories with a net positive (negative) movement along axis X. Motor coated beads (**A** and **E**) displayed a directed movement towards the midline of the pattern (X=0), where they accumulated. Instead, passive beads (**I**) showed a diffusive exploration between the two nucleation lines. **B**, **F** and **J**: Bead position along the axis (*X*) of the actin filaments as a function of time. Most beads take less than 200 s to reach the center of the network. **C**, **G** and **K**: Bead positions in the network 30 min after actin polymerization was initiated. Because they are too small (200-nm diameter) to be clearly visualized, the beads are here represented by 1.2-μm disks, corresponding to a fivefold increase from their actual size. **D**, **H** and **L**: Distribution of bead positions along axis X at steady state (solid line) and when the beads where first detected (dashed line). The nucleation lines are marked by the green shaded areas. The surface fraction of the network occupied by the beads was maximal at the center but remained smaller than 5·10^−4^, corresponding at most to one bead every 60 μm^2^. Because of data folding (see Methods) and bead-size magnification, the surface fraction occupied by the beads in panels **C**, **G** and **K** is 1,512-fold that measured in reality.

As a control, we also studied transport of beads passivated with BSA (Fig. 2I-L). These beads displayed a diffusive exploration of the space between nucleation lines, with trajectories that could start near a nucleation line and later explore regions near the opposite nucleation line (Fig. 2I and J), travelling long distances across the midline of the pattern. Accordingly, the distribution of bead positions at steady state was nearly flat (Fig. 2K and L). We note that passive interactions with the actin network or the substrate nevertheless resulted in a small centering effect, as the steady state distribution was slightly, but significantly, more peaked than the distribution of initial positions (compare dashed and solid lines in Fig. 2L).

### Ensemble analysis of bead transport

To characterize bead transport, we computed the time- and ensemble-averaged squared displacement of the beads—the mean squared displacement MSD(*τ*)—as a function of the lag time *τ*. We recall that MSD(*τ*) = 2*D τ* for a purely diffusive one-dimensional transport characterized by a diffusion coefficient *D*, whereas a purely convective transport of velocity *V* should obey MSD(*τ*) = *V*^2^ *τ*^2^. In the case of beads coated with myosin V, motion analysis along the axis of the actin filaments (axis X; Fig. 1A) revealed that the relation MSD(*τ*) was intermediate between those expected for pure diffusion and convective transport, corresponding to persisting diffusion with MSD(*τ*) ∝ *τ*^1.2^ (*τ* < 50 s; Fig. 3A). This behavior betrayed directed movements driven by myosin-V activity towards the midline of the pattern (Fig. 2A-B). Along the perpendicular axis (axis Y; Fig. 1A), the mean squared displacement was smaller and displayed a sub-diffusive behavior MSD(*τ*) ∝ *τ*^0.8^ (*τ* < 80 s; Fig. 3A).

**Figure 3:**
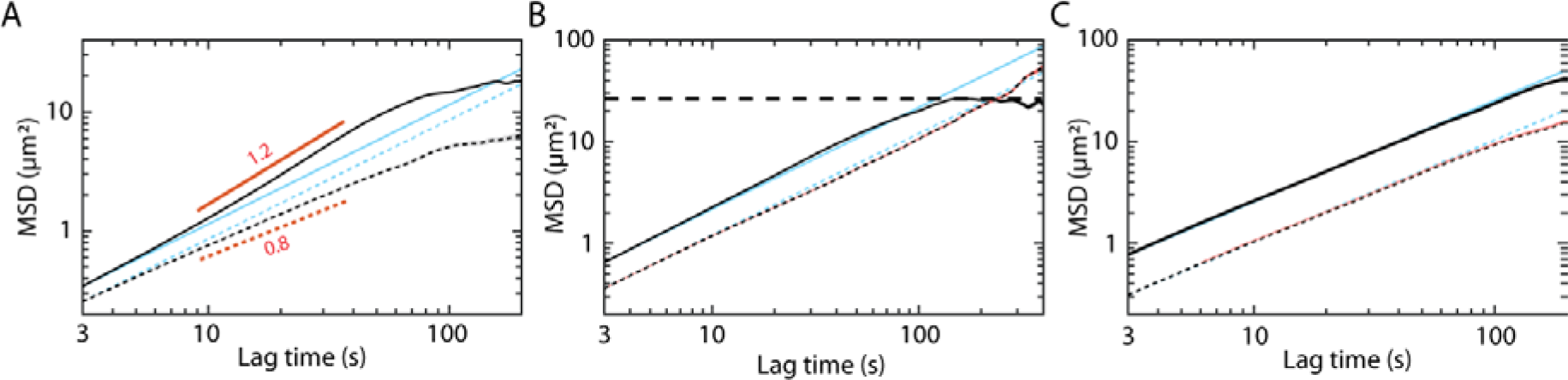
Global analysis of bead transport. The mean squared displacement (MSD) along the X-axis (i.e. parallel to the actin filaments; solid black lines) and along the Y-axis (i.e. perpendicular to the actin filaments; dotted black lines) is plotted as a function of the lag time (*τ*) for myosin-V-coated beads (**A**), for myosin-II-coated beads (**B**), and for BSA-coated beads (**C**). In all cases but for myosin-V coated beads, the relation is well described by MSD(*τ*) = 2*Dτ* for *τ* < 100 s, corresponding to a diffusive one-dimensional transport with a diffusion coefficient *D*. Fits to the data (cyan) yield: *D*_*X*_ = 0.062±0.004 μm^2^/s (*τ* < 10 s) and *D*_*Y*_ = 0.036±0.002 μm^2^/s (n = 205) for myosin-V coated beads, *D*_*X*_ = 0.119±0.001 μm^2^/s and *D*_*Y*_ = 0.054±0.001 μm^2^/s (n= 973) for myosin-II coated beads, and *D*_*X*_ = 0.129±0.002 μm^2^/s and *D*_*Y*_ = 0.052±0.001 μm^2^/s (n = 538) for BSA-coated beads. At long times (*τ* > 100 s), the mean squared displacement of myosin-II coated beads along the X-axis saturates (dashed line), revealing confinement, but not along the Y-axis. Along the X-axis, the relation MSD(*τ*) for myosin-V coated beads is superlinear for *τ* ≤ 50 s, corresponding to persistent diffusion with MSD(*τ*) ∝ *τ*^1.23^ (power-law fit for *τ* ≤ 50 s with R^2^ = 0.9998; [1.21, 1.25], 95% confidence bounds; red solid line), and shows signs of saturation at larger lag times. This behavior is associated with sub-diffusion along the Y-axis according to MSD(*τ*) ∝ *τ*^0.81^ (power-law fit for *τ* ≤ 80 s with R^2^ = 0.9991; [0.79, 0.82], 95% confidence bounds; red dotted line). For each value *τ* of the lag time, the MSD was first time-averaged along each bead trajectory of longer duration than *τ* and then ensemble-averaged over all the trajectories that were detected.

In contrast, with beads coated with myosin II and for lag times *τ*< 100s, we observed that the bead ensemble had a transport behavior dominated by free diffusion (Fig. 3B). The diffusion coefficient was about 2.5 larger along the X-axis than along the Y-axis, reflecting the interaction of the beads with the anisotropic actin network. Beads near the nucleation lines displayed clear directed movements towards the center of the pattern (Fig. 2E and F), as also observed with myosin V. However, although there were signs of persistent diffusion along the X-axis and of sub-diffusion along the Y-axis, as observed with myosin-V-coated beads, the signatures of directed movements were weak in our ensemble analysis of myosin-II-coated beads. At long times (*τ*> 100s), the mean squared displacement along the X-axis clearly saturated to a constant value: the beads were effectively trapped; confinement was not observed along the Y-axis, resulting in larger mean squared displacements along this axis than along the X-axis. With beads passivated by BSA, the mean squared displacement displayed anisotropic diffusion resembling that observed with myosin-II-coated beads, but with the important difference that no confinement was observed at long times for passive beads (Fig. 3C). In return, the saturation of the mean squared displacement that we observed with myosin-II-coated beads betrayed the activity of the motors, which strived to maintain the beads near the center of the network.

### Spatial diffusion and drift profiles

Because the density and net polarity of the actin network varied with position along the X-axis, the transport properties of the beads were expected to be inhomogeneous. Spatial heterogeneities were averaged in the calculation of the mean-squared displacements (Fig. 3), which corresponded to ensemble averages of bead displacements, irrespective of their position in the network. To reveal the effects of the actin architecture on transport, we determined the local diffusion coefficient *D*(*X*) and drift velocity *V*(*X*) of the beads as a function of position *X* (Methods; Fig. 4). We observed that the drift velocity of the beads varied like the net polarity of the network (Fig. 1C, inset): the bead velocity saturated to a maximal (absolute) value near the nucleation line and declined (in absolute value) as the beads approached the center, changing sign there (Fig. 4A and D). In addition, the local diffusion coefficient displayed a local minimum near the center, where there was less actin (Fig. 4B and E).

**Figure 4:**
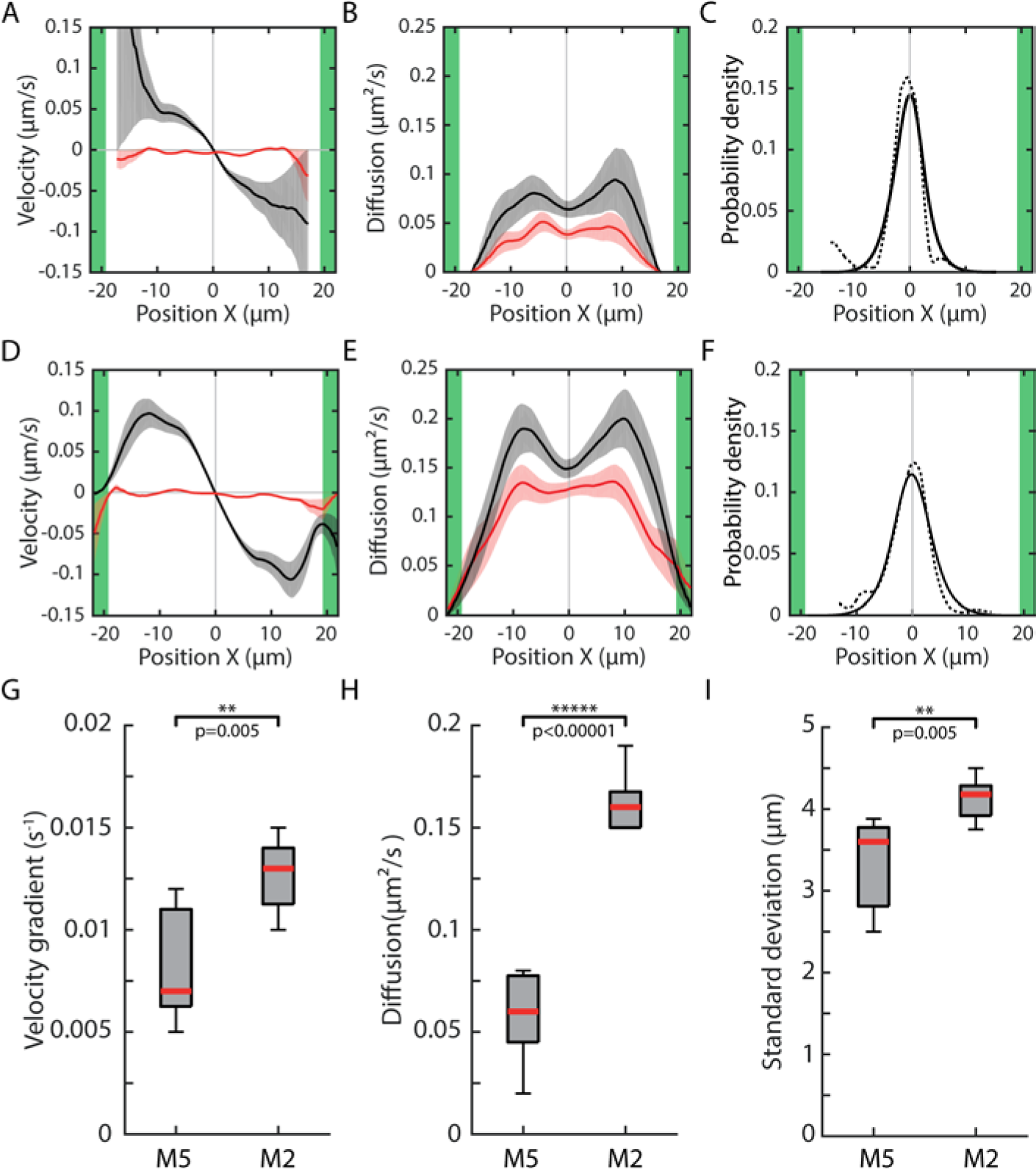
Local analysis of active bead transport: velocity and diffusion-coefficient profiles. For myosin-V-coated beads (**A-C**) and myosin-II-coated beads (**D-F**), we plot the velocity (**A** and **D**) and diffusion coefficient (**B** and **E**) as a function of position *X* along the axis of the actin filaments (black lines: along X axis; red lines: along Y axis), as well as the steady-state distributions of bead positions (**C** and **F**) that were measured directly (dashed lines; same data as Fig. 2D and H) and predicted by a diffusion-drift process (solid lines; see Methods). In **G**-**I**, we show box plots of velocity gradient at the center (**G**), of the diffusion coefficient at the center (**H**) and of the standard deviation of bead position (**I**) for the two types of motor-coated-beads. The data shown in **G-I** results from N = 7 experiments in which myosin-V-coated beads and myosin-II-coated beads moved in the same antiparallel actin network, allowing for a direct comparison of their transport properties; statistical significance (stars) was assayed by paired-sample t-tests with p-values indicated on the figure.

Remarkably, the measured steady-state distributions of bead position (dashed lines in Fig. 4C and F) was well described by that predicted from a diffusion-drift process (*P*_*DD*_) with the measured velocity and diffusion profiles (solid lines in Fig. 4C and F). In a diffusion-drift process, a convective flux *J*_*C*_ = *P*_*DD*_ *V* transporting the beads towards the center of the network competes with a diffusive flux *J*_*D*_ = −*D dP*_*DD*_/*dX* that homogenizes the bead distribution. At steady state, because there is no apparent collective directional movement, the total flux *J*_*C*_ + *J_D_* = 0, which yields 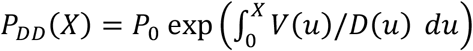. Because *V*(*X*) is odd and *D*(*X*) is even, the distribution *P*_*DD*_ (*X*) shows a peak at *X* = 0. The variance of bead position associated with a Gaussian approximation to the peak is given by 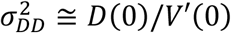, where the prime corresponds to a spatial derivative. Thus, the steeper the velocity gradient and the lower the diffusion, the narrower the distribution. Notably, as the result of their larger speeds, myosin-II-coated beads displayed steeper gradients *V*′(0) than myosin-V-coated beads (Fig. 4G). However, diffusion coefficients *D*(0) were also much larger with myosin II than with myosin V (Fig. 4H). The standard deviation σ of bead position characterized the precision of bead positioning. We found *σ*_MV_ = 3.3 ± 0.6 μm with myosin V and *σ*_MII_ = 4.1 ± 0.3 μm with myosin II (mean±SEM; n = 7 experiments; Fig. 4I). Thus, active bead positioning at the center of the network was more precise with myosin V, by 25%. Note that these standard deviations are significantly smaller, by a factor two or more, than the characteristic length *λ* of the net-polarity gradient. Thus, bead positioning by motors was more precise than the lengthscale that characterizes structural heterogeneities of the actin network.

### 3-state model of bead transport

To understand the physical origin of active bead positioning in antiparallel filament networks, we worked within the framework of a 3-state model (6). A bead could be in one of the following three states (Fig. 5A): (state 1) attached to a filament oriented in the positive direction and moving along this filament at velocity +*ν*, (state 2) attached to a filament oriented in the negative direction and moving along this filament at velocity − *ν*, or (state 3) detached and freely diffusing with a diffusion coefficient *D*_∅_. We considered a uniform detachment rate constant *k_OFF_* from the attached states 1 and 2 to the detached state 3. In contrast, attachment rate constants depended on the local density *ρ*^±^(*X*) of filaments of any given polarity: *k*_3→1_(*X*) = *k_ON_ ρ*^+^(*X*) and *k*_3→2_(*X*) = *k_ON_ ρ*^−^(*X*). Thus in the model, beads are more likely to attach to a filament of a given polarity when the corresponding filament density is larger than the density of filaments with the opposite polarity; this ingredient is key, because it provides a mechanism for sensing the net polarity of the network. Within this framework, we define two characteristic timescales: the mean time *τ_ON_* = 1/*k_OFF_* spent in an attached state (state 1 or 2) and the mean time *τ_OFF_* = 1/(*k_ON_ ρ*_0_) spent in the detached state (state 3), were *ρ*_0_ = *ρ*^+^(0) + *ρ*^−^(0) is the total actin density at the center (*X* = 0) of the network. Note that the mean time spent in the detached state depends on the total actin concentration and thus on position in the network; we use here as a reference the value of this characteristic time at the center, *τ_OFF_*. We also define two characteristic lengthscales: the mean distance *ℓ*_*B*_ = *vτ_ON_* by which a bead travels along a filament in an attached state—the *ballistic length*—and the distance 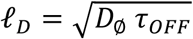 that controls the mean-squared displacement of a bead in the detached state as a result of Brownian motion—the *diffusion length*. In practice, we used exponential density profiles *ρ*^±^(*X*) = (*ρ*_0_/2) exp(∓*X*/*λ*), as observed in experiments (Fig. 1C).

**Figure 5:**
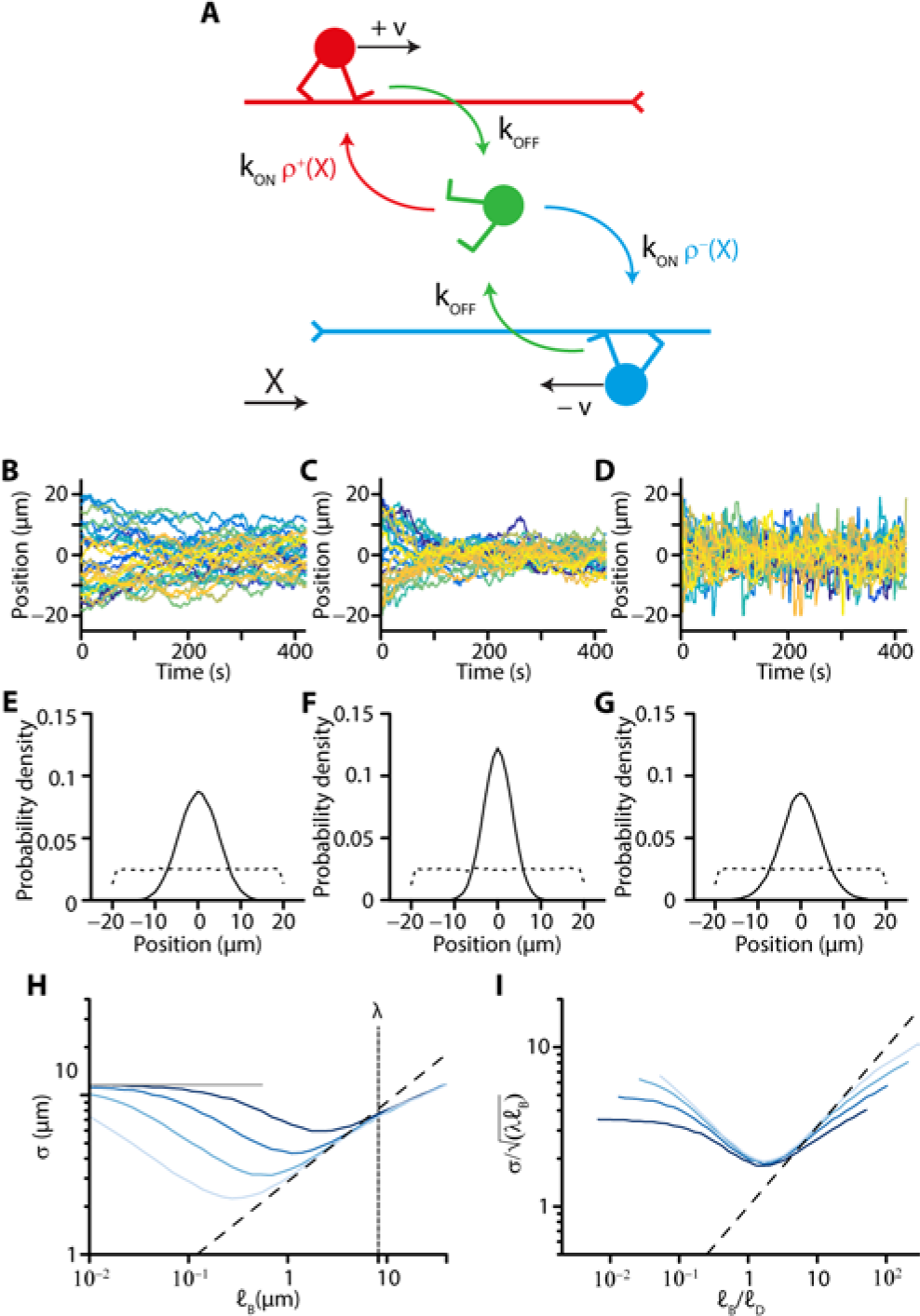
Active positioning in a 3-state model of bead transport. **A:** Model myosin-coated beads can be attached to actin filaments and moving towards their barbed ends at velocity +*v* (red; state 1) or − *v* (blue; state 2) or detached and freely diffusing with a diffusion coefficient *D*_∅_ (green; state 3). Attached beads detach at a rate *k*_OFF_ that does not depend on position in the network nor on filament polarity. Detached beads at position *X* can attach to filaments of a given polarity (+ or −) at rates *k*_ON_*ρ*^±^(*X*) proportional to the local density *ρ*^±^(*X*) = *ρ*^±^(0) exp(∓ *X*/**λ**) of the filaments, where **λ** corresponds to the mean filament length. **B-D:** Model-bead trajectories at low (*v* = 0.5 μm/s and *ℓ*_*B*_/*ℓ*_*D*_ = 0.25; **B**), intermediate (*v* = 2 μm/s and *ℓ*_*B*_/*ℓ*_*D*_ = 1; **C**) and high (*v* = 8 μm/s and *ℓ*_*B*_/*ℓ*_*D*_ = 4; **D**) velocity, which corresponds to increasing values of the ratio *ℓ*_*B*_/*ℓ*_*D*_. Other parameter values in SI Appendix, Table S2 (Case 1). **E-G:** Model-bead distribution at the start of the simulation (dotted line) and at steady state (solid line) corresponding to the simulated trajectories shown in **B-D**, respectively. **H:** Standard deviation σ of bead position at steady state as a function of the ballistic length *ℓ*_B_ = *v*/*k*_OFF_ for four different values (0.3, 0.6, 1.1, and 2.2 μm, from light to dark blue) of the diffusion length 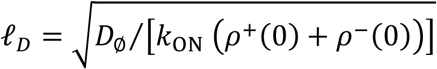 at the center (*X* = 0) of the filament network. The ballistic length was varied by changing the value of *v*; the diffusion length was varied by changing the value of *D*_∅_; other parameters in SI Appendix, Table S2 (Case 1). **I:** Same data as in **H**, but using normalized coordinates. In **H-I**, the dashed line corresponds to the relation 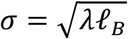.; in **H**, the horizontal grey solid line corresponds to the value σ = *L*^2^/12 expected for free one-dimensional diffusion in a box of size *L* = 40 μm.

We simulated the stochastic dynamics of beads—called ‘model beads’ in the following (SI Appendix); at the start of the simulations, the model beads were randomly distributed in the antiparallel network. With two lengthscales and two timescales, we could operate in four distinct regimes of motion (SI Appendix, Fig. S3). In the regime of high attachment rates (*τ*_*ON*_ ≫ *τ*_*OFF*_), model beads are expected to move at velocity ±*v* near the edges of the antiparallel network, where |*X*| ≅ *L*/2 ≫ *λ* and the net polarity Φ(*X*) ≅ ±1. However, the ensemble-averaged bead velocities that were measured at these positions in the antiparallel network (Fig. 4A and D) or at all positions in parallel networks (SI Appendix, Fig. S1) were almost one order of magnitude smaller than the natural velocity of the motors, as estimated in standard gliding assays (1). To account for this observation, we had to ensure in our simulations that the attachment probability of a bead was low, on the order of 10%, and thus choose *τ*_*ON*_ ≪ *τ*_*OFF*_. With this choice, we were able to reproduce (Fig. 5C) the active funneling observed with myosin-coated beads (Fig. 2B and F) when the model beads in our simulations traveled comparable distances in the attached and detached states (*ℓ*_*B*_~*ℓ*_*D*_). In this regime, the steady-state distribution of the model beads was peaked at the center of the network (Fig. 5C and F). Strikingly, by continuously increasing *ℓ*_*B*_ while keeping *ℓ*_*D*_ at a fixed value, we found that the standard deviation of model-bead position at steady state first decreased and reached a minimum at *ℓ*_*B*_ ≅ 1.65 *ℓ*_*D*_, corresponding to an optimum of centering precision (Fig. 5H-I), before increasing again.

A hierarchy of lengthscales emerged naturally in the model: *L* > *λ* > *ℓ*_*B*_~*ℓ*_*D*_. The mean-actin length *λ*, which sets the lengthscale of the net-polarity gradient of the antiparallel network, had to be smaller than the size *L* of the network to ensure that the network was structurally inhomogeneous. Importantly, the condition *λ* > *ℓ*_*B*_ ensured that the bead trajectories in the attached state were short enough to sample the net-polarity gradient from detachment-reattachment events. Finally, centering required that the beads did not diffuse too far in the detached state (*ℓ*_*B*_~*ℓ*_*D*_; Fig. 5 and SI Appendix, Fig. S4) in order to keep a directionality of motion towards the center.

Provided that the density of the network varies slowly with position (*λ* ≫ *ℓ*_*B*_, *ℓ*_*D*_), we could calculate analytically an effective drift velocity *v*_*EFF*_ (*X*) and an effective diffusion coefficient *D*_*EFF*_ (*X*) for the model beads (SI Appendix, section 3). In the experimentally-relevant limit of frequent detachment (*α* = *τ_ON_*/*τ_OFF_* ≪ 1), we found *v*_*EFF*_ (*X*) ≅ −*v α* sinh(*X*/*λ*) and 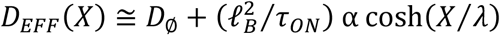. Notably, over a broad range of parameter values (0.1 < *ℓ*_*B*_/*ℓ*_*D*_ < 10; SI Appendix, Fig. S6), the steady-state distribution of model-bead position was well approximated by that given by a diffusion-drift process using these velocity and diffusion profiles: 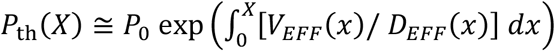. As in experiments, this distribution peaked in the center (*X* = 0): the active transport process brought the beads, on average, to the position where the net polarity of the network vanishes. Using a Gaussian approximation of the peak (SI Appendix, Eq S13), we estimate the variance of model-bead position 

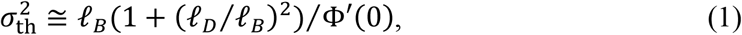

where the polarity gradient at the center Φ′(0) = 1/*λ* is given by the mean filament length. In qualitative agreement with numerical estimates (Fig. 5H-I), the relation between the variance 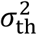 and the ballistic length *ℓ*_*B*_ (Eq. 1) shows a minimum; the minimal variance occurs here at *ℓ*_*B*_ = *ℓ*_*D*_, where 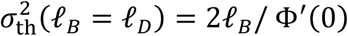. Clearly, at fixed *ℓ*_*B*_, 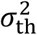 is minimal at *ℓ*_*D*_ = 0.

By fitting the theory to our experiments, we were able to get numerical values for the parameters in the model. The actual value *ℓ*_*D*_/*ℓ*_*B*_ is uncertain but our theory indicates that this ratio must display an upper bound (~4 for myosin V, ~1.5 for myosin II; see SI Appendix, section 3) to account for the observed minimum of the effective diffusion coefficient at the center of the network (Fig. 4B and E; SI Appendix Fig. S5D). In the limit of small *ℓ*_*D*_, parameter values are listed in Table 1. In particular, we inferred the ballistic length *ℓ*_*B*_ = σ^2^/*λ* from the measured values of the variance σ of bead position (Fig. 4I) and of the mean actin length *λ* (Fig. 1B). We found *ℓ*_*B*_ = 0.77 ± 0.39 μm (n = 11) for myosin V that was smaller than the value *ℓ*_*B*_ = 1.25 ± 0.47 μm (n = 16) for myosin II, as required by Equation 1 (with *ℓ*_*D*_/*ℓ*_*B*_ ≅ 0) to ensure that bead positioning in a given actin network was more precise for with myosin V (Fig. 4). Near the optimal precision of positioning (*ℓ*_*D*_/*ℓ*_*B*_ ≅ 1), the ballistic length is half the value estimated when passive diffusion is ignored (*ℓ*_*D*_/*ℓ*_*B*_ ≅ 0).

**Table 1:**
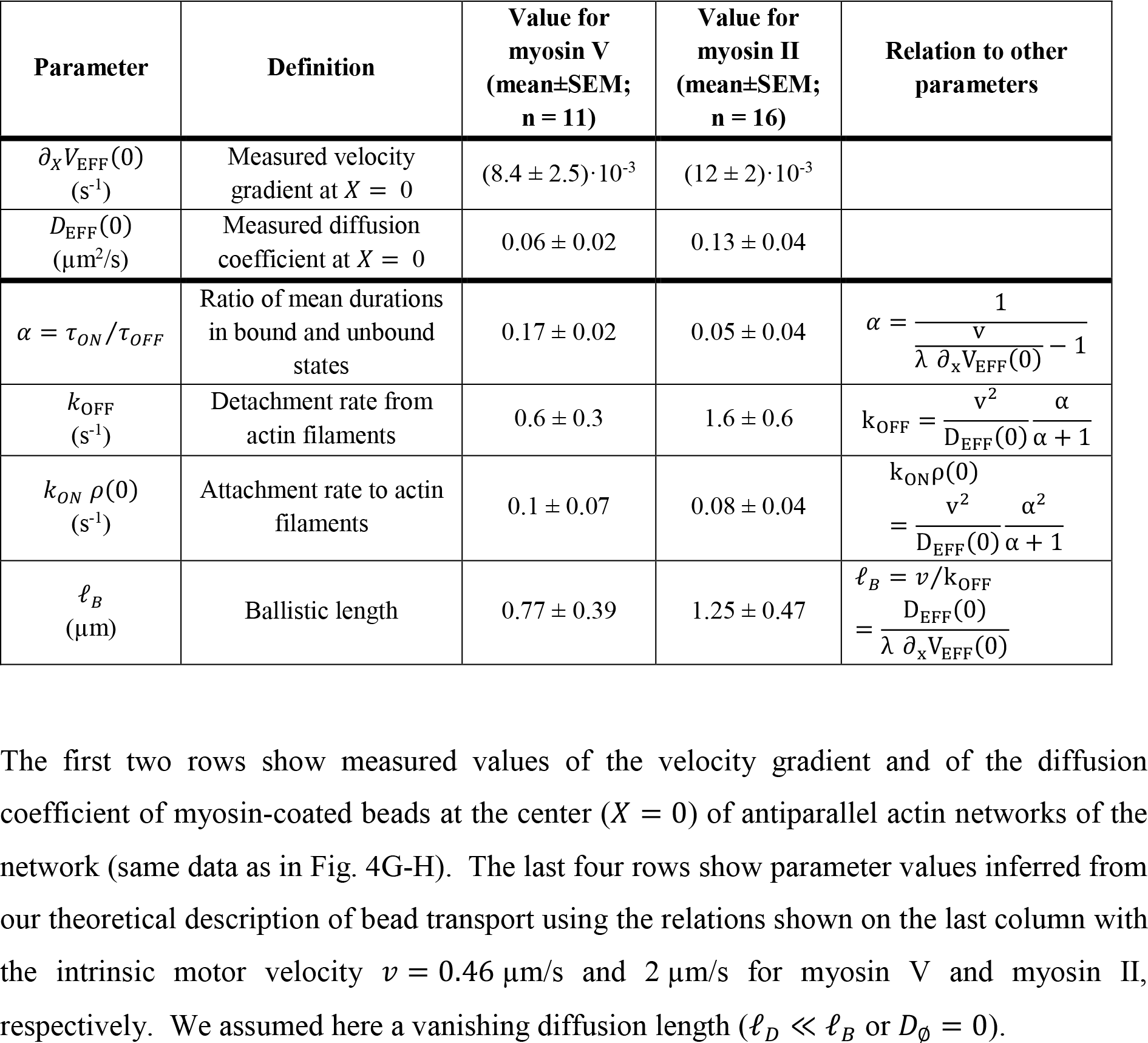
parameter values. The first two rows show measured values of the velocity gradient and of the diffusion coefficient of myosin-coated beads at the center (*X* = 0) of antiparallel actin networks of the network (same data as in Fig. 4G-H). The last four rows show parameter values inferred from our theoretical description of bead transport using the relations shown on the last column with the intrinsic motor velocity *v* = 0.46 μm/s and 2 μm/s for myosin V and myosin II, respectively. We assumed here a vanishing diffusion length (*ℓ*_*D*_ ≪ *ℓ*_*B*_ or *D*_∅_ = 0).

Irrespective of the actual value of *ℓ*_*D*_/*ℓ*_*B*_, the model predicts that the precision of bead positioning (Eq. 1) should not depend on the size (*L*) of the actin network. This prediction was actually confirmed experimentally (SI Appendix, Fig. S2A). In addition, the standard deviation σ of bead position is expected to depend on the total actin density *ρ*_0_ at the center, as can be seen by rewriting Equation 1 as 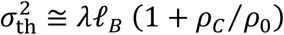, in which 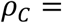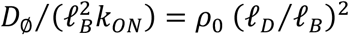 corresponds to the critical actin density at which *ℓ*_*D*_ = *ℓ*_*B*_. In the regime where the actin density is low enough (*ρ*_0_ ≪ *ρ*_*C*_), the variance of bead position ought to be inversely proportional to the actin density *ρ*_0_. Conversely at high actin densities (*ρ*_0_ ≥ *ρ*_*C*_), the positioning precision should only weakly depend on the total actin density *ρ*_0_. Increasing the concentration of monomeric actin in the polymerization mix from 2 μM to 5 μM had no significant effect on the measured value of σ (SI Appendix, Fig. S2B), suggesting that the condition *ρ*_0_ ≥ *ρ*_*C*_ was satisfied and therefore that *ℓ*_*B*_ ≥ *ℓ*_*D*_ in our experiments.

## Discussion

In this paper, we have presented a minimal in-vitro assay to study active cargo transport in mixed-polarity networks of cytoskeletal filaments. Focusing on the acto-myosin system, we found that myosin-coated beads can sense the net polarity of the network, which provides a cue for directed movements. Accordingly, the velocity of bead motion varied with the net actin polarity and went to zero at positions where the polarity vanished (Inset of Fig. 1C, Fig. 4A and D). As a result, the beads were actively trapped at these positions, providing a general mechanism of cargo positioning in mixed-polarity networks that does not depend on specific molecular recognition between a cargo and its target (21).

At steady state, the myosin-coated beads were confined within a region (Fig. 4) that was significantly smaller than the characteristic lengthscale associated with the net-polarity gradient of the antiparallel network, which was here set by the mean actin length *λ* (Fig. 1). Theoretical analysis via a coarse-grained model revealed that positioning depends not only on the steepness of the net-polarity gradient, but also on an interplay between the ballistic length *ℓ*_*B*_ and the diffusive length *ℓ*_*D*_ of the myosin-coated beads (Eq. 1). The ballistic length corresponds to the mean length that the myosin-coated beads travel in an attached state to actin before detaching and in turn probe the net polarity of the network. As long as *ℓ*_*B*_ > *ℓ*_*D*_, the shorter the ballistic length, the finer the sampling of the net-polarity profile of the network, and thus the more precise the positioning (Fig. 5). However, the ballistic length cannot be too short for precise positioning: diffusive transport becomes limiting when *ℓ*_*B*_ < *ℓ*_*D*_. Our model suggests that precise positioning emerges because the ballistic length *ℓ*_*B*_ (~1μ*m*) is shorter than the mean-actin length *λ* (~8 μ*m*). Within our theoretical framework, active transport is described as a stochastic sequence of processive runs and diffusive searches that results in biased random walk towards a direction dictated by the net-polarity gradient.

An even simpler theoretical approach had been previously developed to describe dynein-driven transport in the mixed-polarity network of microtubules found in dendrites (7). However, this earlier work ignored diffusion in a detached state of the cargo and thus cannot describe the condition for optimal bead positioning that we reveal here (Fig. 5H-I). Our description extends a theory developed originally to capture endosomal transport during the asymmetric division of sensory organ precursors in Drosophila (6); we here consider slowly varying density profiles of antiparallel filament networks instead of overlapping homogeneous networks of opposite polarities.

Excluded-volume interactions between motor particles can lead to traffic jams, which shape the particle distribution at steady state (22). However, in our experiments, the surface fraction of the network occupied by the myosin-coated beads was so low (e.g. ~10^−4^ for the myosin-II coated beads shown in Fig. 2G) that we could safely neglect steric bead interactions. Unidirectional transport has also been extensively studied in the context of cylindrical organelles such as filopodia or stereocilia. There, predicted motor distributions are critically controlled by boundary conditions, due to the influx of motors at the open end of the half-closed tube describing the geometry of those systems (23–25). In our experiments, beads instead enter the network by sedimentation from the bulk, in a direction perpendicular to the actin sheet. The sedimentation flux is uniform and very small (e.g. 0.025 bead/sec distributed over an area 40 μm × 350 μm of the actin network in the experiment corresponding to data shown in Fig. 2F, G and H). In addition, the width of the experimental bead distribution is one order of magnitude smaller than the width of the actin network (*L* = 40μm) and is vanishingly small at the boundaries (nucleation lines; Fig. 2). Thus at the edges, the bead influx along the axis of the filaments must be small.

Myosin-coated beads could in principle interact with multiple filaments of mixed polarity, resulting in a tug-of-war (26, 27). However, a tug-of-war is inherently unstable, resulting in bi-directional movements and bead switching between filaments (5, 13). Although we did not explicitly account for bead interaction with multiple filaments in our description of transport (SI Appendix), we observed that positioning was more precise in our simulations (SI Appendix, Fig. S7) when the beads were allowed to switch directly between the two attached states (States 1 and 2; Fig. 5A). In this case, analytical calculations in the small-diffusion limit (*L* > **λ** > *ℓ*_*B*_ ≫ *ℓ*_*D*_) revealed that the variance of the steady-state distribution of bead position was also given by Equation 1, albeit with a renormalized ballistic length 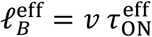 (SI Appendix). With direct filament switching, the ballistic length was reduced. This is because the effective mean time 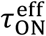 of a run in a given direction when the beads are attached to the filaments—the *persistence* time—is reduced by direct filament switching. Note that in the absence of switching, the persistence time is simply given by the mean attachment time 1/*k*_off_. The beads may also sense the net polarity of the network by switching only, i.e. without ever detaching from the filaments, and thus position themselves at the center of the antiparallel network (SI Appendix, Fig. S8). However, this mechanism would produce much steeper velocity gradients that those reported here and the diffusion profile would display a maximum at the center, in contradistinction to our observations (Fig. 4B and E; SI Appendix, Fig. S9).

We studied active transport of beads coated either with myosin Va or skeletal heavy-mero myosin II. Both types of motors are double-headed molecular motors with movements directed towards the barbed ends of the actin filaments, thus away from the nucleation lines of actin polymerization (Fig. 1). However, their biophysical properties, as well as their functions in vivo, differ strongly (28). Myosin V is a processive molecular motor involved in intracellular transport (29), whereas skeletal myosin II is a non-processive motor involved in muscle contraction, not in cargo transport. Yet, both types of myosin-coated beads displayed similar transport behaviors down the net-polarity gradients of antiparallel actin networks, resulting in bead positioning (Fig. 2). Our data adds to the available evidence (30–32) that non-processive motor molecules, here myosin II, can be turned into a processive transporter if the motors work as a group on the cargo.

Positioning was more precise with beads coated with processive myosin-V than with non-processive myosin-II motors. Local analysis of bead velocity and diffusion revealed that the positioning could be described by a diffusion-drift process, for which the standard deviation of bead position is set by the ratio of an effective diffusion coefficient and an effective velocity gradient (Fig. 4). The velocity gradient was larger by a factor ~1.5 with myosin II than with myosin V. However, the effective diffusion coefficient was larger by a bigger factor (~2.2) with myosin II, resulting in less precise cargo positioning with this motor type. Our 3-state model of active bead transport reveals that the key motor property that controls positioning precision (Eq. 1) is actually the ballistic length *ℓ*_*B*_ = *v* /*k*_OFF_, which is given by the ratio of the bead velocity *v* in an attached state to actin and the bead detachment rate *k*_OFF_.

Although myosin-II and myosin-V motors have been extensively characterized at the single molecule level (28), beads are each transported by a group of motors. How multiple motor molecules coordinate or impede their movements to collectively mediate cargo motion remains unclear (11, 33). Relating the stochastic properties of bead dynamics to known properties of single motors is an avenue for future research but is beyond the scope of this work. Generally, increasing the number of motors on a bead is expected to increase the ballistic length *ℓ*_*B*_ (32). As already discussed above, whether or not increasing *ℓ*_*B*_ results in more precise positioning actually depends on how the ballistic length compares with the diffusive length *ℓ*_*D*_ (Fig. 5).

Our analysis of active transport indicates that bead positioning not only depends on intrinsic motor properties but also on the architecture of the actin network (Eq. 1). In particular, the lower the mean actin length **λ**, and thus the steeper the net-polarity gradient (Fig. 1C, inset), the sharper the positioning of the beads at the center of the antiparallel network. This condition holds true as long as the ballistic length *ℓ*_*B*_ remains smaller than **λ** to ensure centering (i.e. σ < **λ**). In addition, the precision of bead positioning can depend on the total actin density, but only at densities that are low enough that 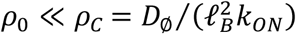. In this case, the higher the actin density, the more precise the positioning. At high actin densities (*ρ*_0_ ≥ *ρ*_*C*_), positioning instead becomes independent on the total actin density, as observed in our experiments (SI Appendix, Fig. S2B). Altogether, our results emphasize the interplay between intrinsic motor properties and the architecture of the actin network for active positioning of myosin-transported cargoes.

## Materials and Methods

### Micro-patterning of an actin nucleation-promoting factor (pWA)

Details of the micro-patterning protocol have been published elsewhere (18, 19). In short, glass coverslips were oxidized with an oxygen plasma for 3 min and incubated with 0.1 mg/ml Poly(L-lysine)-graft-poly(ethylene glycol) (PLL-g-PEG; Jenkem Technology) in 10 mM HEPES at pH = 7.4 for 30 min. The passivated surface was exposed to deep UV light (wavelength: 180 nm; UVO Cleaner Unit 342, Jelight Company INC) for 5 min through a transparent micropattern that was drawn on a chromium synthetic-quartz photomask (Toppan Photomasks). The coverslips were then incubated with 1 μM pWA in a buffer containing 50 mM KCl, 1 mM MgCl2, 1 mM EGTA and 10 mM imidazole-HCl (pH = 7.8) for 10 min. In our experiments, the pattern was typically composed of seven parallel lines that were 3-μm wide, 350-μm long and spaced by 40 μm.

### Actin polymerization

To induce actin polymerization from surface micro-patterns of the nucleation-promoting factor pWA, 4 μM globular actin (Tebu-Bio), 12μM profilin and 10 nM Arp2/3 complex (Tebu-Bio) were mixed in a buffer containing 10 mM imidazole-HCl (pH = 7.8), 50 mM KCl, 1 mM MgCl_2_, 2.6 mM Na_2_ATP, 56 mM dithiothreitol, 0.1 mg/ml glucose, 3.7 Units/ml catalase, 37.3 Units/ml glucose oxidase and 0.4% (w/w; macrocospic viscosity 800 cP) methylcellulose. To visualize actin filaments, 20% of the monomers were labelled with a fluorophore (Alexa568; Life Technologies). A Peltier element and feedback control (Warner Instruments) ensured that experiments were performed at a temperature of 27°C. A surface micropattern of seven parallel actin-nucleation lines resulted in six identical antiparallel networks of overlapping actin filaments.

### Bead functionalization with myosins

To study cargo transport in actin networks, we added functionalized beads to the actin-polymerization solution described in the preceding section. We used polystyrene beads of 200 nm in diameter (Life Technologies). A 1-μl volume of a bead stock at 0.1% (w/w), corresponding to ~2.3×10^8^ beads, was added to 24 μl of a buffer containing 80 mM KCl, 10 μM EDTA, 1 mM EGTA and 10 mM imidazole-HCl. The bead suspension had a concentration of 40 μg/ml. The beads were washed by centrifuging the solution at 20,000 g for 30 min at 4°C and by then replacing 19 μl of the supernatant with the same volume of buffer. After dispersing the beads in the solution by manual trituration followed by sonication during 30 s, we added 3 μl of a myosin solution at 0.44 μM and allowed the myosin molecules to absorb onto the surface of the beads for an incubation time of 10 min. Finally, a volume of 3 μl of BSA at 10% (w/w) was added to passivate the portion of the beads’ surface that was not occupied by the myosin. The initial concentration of the myosin in the solution was 47 nM. For bead-transport studies, the concentration of functionalized beads was adjusted by dilution to ~6×10^5^ μl^−1^.

We used either of two types of myosin molecules. First, recombinant double-headed myosin Va missing the C-terminal globular tail were produced as described (34). Second, heavy-mero myosin II, purified from rabbit pectoral muscle, was kindly provided by Matthias Rief’s group (TUM, Germany). In control experiments with passive beads, we omitted to add myosin and instead fully passivated the bead surface with BSA.

### Microscopic observation

Our samples were viewed through a ×20 objective (NA=0.75) of a spinning-disk confocal microscope (Eclipse Ti, Nikon); this low magnification allowed us to record hundreds to thousands of bead trajectories. We recorded time-lapse videos with a CCD camera (CoolSNAp HQ2, Photometrics) at a framerate of 3 s. In the sample plane, the pixel size of the camera was 322.5 nm × 322.5 nm. Time-lapse recordings started 10 min after the injection of the polymerization mix into the flow chamber and lasted for 20 min. Note that the functionalized beads were already present in the solution used for actin polymerization at the initiation of the polymerization process. For the characterization of the actin-network florescence profiles (Fig. 1), we used a ×40 objective (NA=0.75).

### Single-particle tracking

We automatically measured bead trajectories using TrackMate (35), a single-particle tracker developed as a plugin for the image-processing software Image J (National Institute of Health, Bethesda, USA). Bead tracking was performed with a time resolution of 3 s, corresponding to the framerate of our time-lapse videos. An experiment produced up to 5,000 trajectories. We filtered the data in two ways. First, we cut the portions of any trajectory for which the standard deviation of the bead position remained below 0.1 μm during at least 45 s. Some beads took several minutes to start moving after they appeared in the field of view, or showed long pauses in their trajectory before moving again, or stopped until the end of the video. Any given trajectories could thus be parsed in 2‒5 tracks per trajectory during a 20-min recording. Second, we retained only the tracks that explored a region that could be inscribed in a disk with a diameter larger than 2 μm, thus rejecting shorter tracks. This procedure rejected about half of the available tracks. Tracks were typically collected from six identical lanes of the surface micropattern (see above), corresponding each to an area 350 μm × 40 μm. We assumed translational invariance of the actin network in a direction parallel to the nucleation lines as well as periodicity in the perpendicular direction (defined respectively as Y- and X-axis, see Fig. 1). These features allowed for data folding as well as spatial averaging.

In our experiments, there was a slow continuous flux of beads from the bulk to the plane of observation within the actin network. Different beads thus appeared at different times on the actin network. Any given frame of a time-lapse video in turn showed a mix of ‘old’ beads that were given enough time to reach steady state and of ‘young’ beads, mainly positioned near the nucleation lines, which were en-route towards the center of the antiparallel actin network (Fig. 2). To estimate the steady-state distribution of bead positions, we scanned all the tracks of the mobile beads to select those (~100 tracks) that lasted at least 200 s; each bead position *X*(*t* = *T*_EQ_) was then registered at time *t* = *T*_EQ_ = 200 s after the start of its track and we computed the corresponding distribution *P*_M_(*X*). The duration *T*_EQ_ corresponds to the travel time of a bead moving at a mean velocity 0.1 μm/s from a nucleation line to the center of an antiparallel network if the spacing between nucleation lines is 40 μm. To allow for easy comparison of the bead trajectories along the X-axis (Fig. 2B, F and J), the time origin of each track was reset to the time were the bead first appeared on the time-lapse video.

### Profiles of drift velocity and diffusion coefficient

We computed the profiles of the local drift velocity *V*(*X*) and of the local diffusion coefficient *D*(*X*) of the beads as a function of bead position *X*. To estimate *V*(*X*) and *D*(*X*), we measured the distribution of displacements Δ*X*_*i*_(*X*) during a fixed time-interval *τ* of all the beads *i* = 1, …, *N*_*X*_ that are found during their trajectory at a given position *X* with a precision of 0.1 μm and computed the mean value 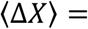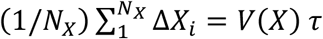 and its variance 〈(Δ*X*)^2^ − 〈Δ*X*〉^2^〉 = 2*D*(*X*) *τ*. Knowing the profiles *V*(*X*) and *D*(*X*), we calculated the probability density 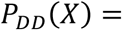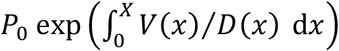 of bead positions expected from a diffusion-drift process at steady state (see main text). In practice, we used *τ*= 12 s; we checked that the estimated probability density *P*_*DD*_(*X*) remained nearly unchanged for variations of *τ* in the range 3-24 s. Over a timescale *τ*, the beads moved on average by less than 100 nm and the net polarity of the actin network (Fig. 1C, inset) varied by less than 2%. The calculated distribution *P*_*DD*_ (*X*) was confronted to the bead distribution *P*_M_(*X*) measured directly by the procedure described in the preceding paragraph (Fig. 4C and F).

## Acknowlegements

We are indebted to M. Rief for providing the heavy-mero myosin II molecules. We acknowledge the use of microscopes from the Cell and Tissue Imaging Center (PICT-IBiSA) of the Institut Curie, which is a member of the French National Research Infrastructure France-BioImaging (ANR10-INBS-04). We thank John Manzi from the Biochemisty, Molecular Biology and Cells platform of Laboratoire Physico-Chimie Curie (Institut Curie) for protein purification and characterization. We also thank Julie Plastino and Cécile Sykes for fruitful discussions and for sharing their expertise about actin polymerization. This work was supported by the French National Agency for Research (ANR-12-BSV5 0014; awarded jointly to P.M. and L.B.), by the Labex Celtisphybio ANR-10-LABX-0038 part of the Idex PSL, by the United States National Institutes of Health R01 Grant GM097348 (awarded to E. M. D. L. C.), and by the European Research Council (741773 (AAA); awarded to L.B.). E.M.D.L.C. was a Mayent-Rothschild Senior Researcher Fellow at the Institut Curie. M.R. is alumnus of the Frontiers in Life Science PhD program of Université Paris Descartes.

## Supporting Online Appendix

Figs. S1 and S2 (section 1), Supplementary Text (sections 2 to 4, with Figs. S3–S9), Table S1–S2 (section 5).

## SI Appendix

### Section 1: Supplementary data (Figs. S1 and S2)

**FIG. S1:**
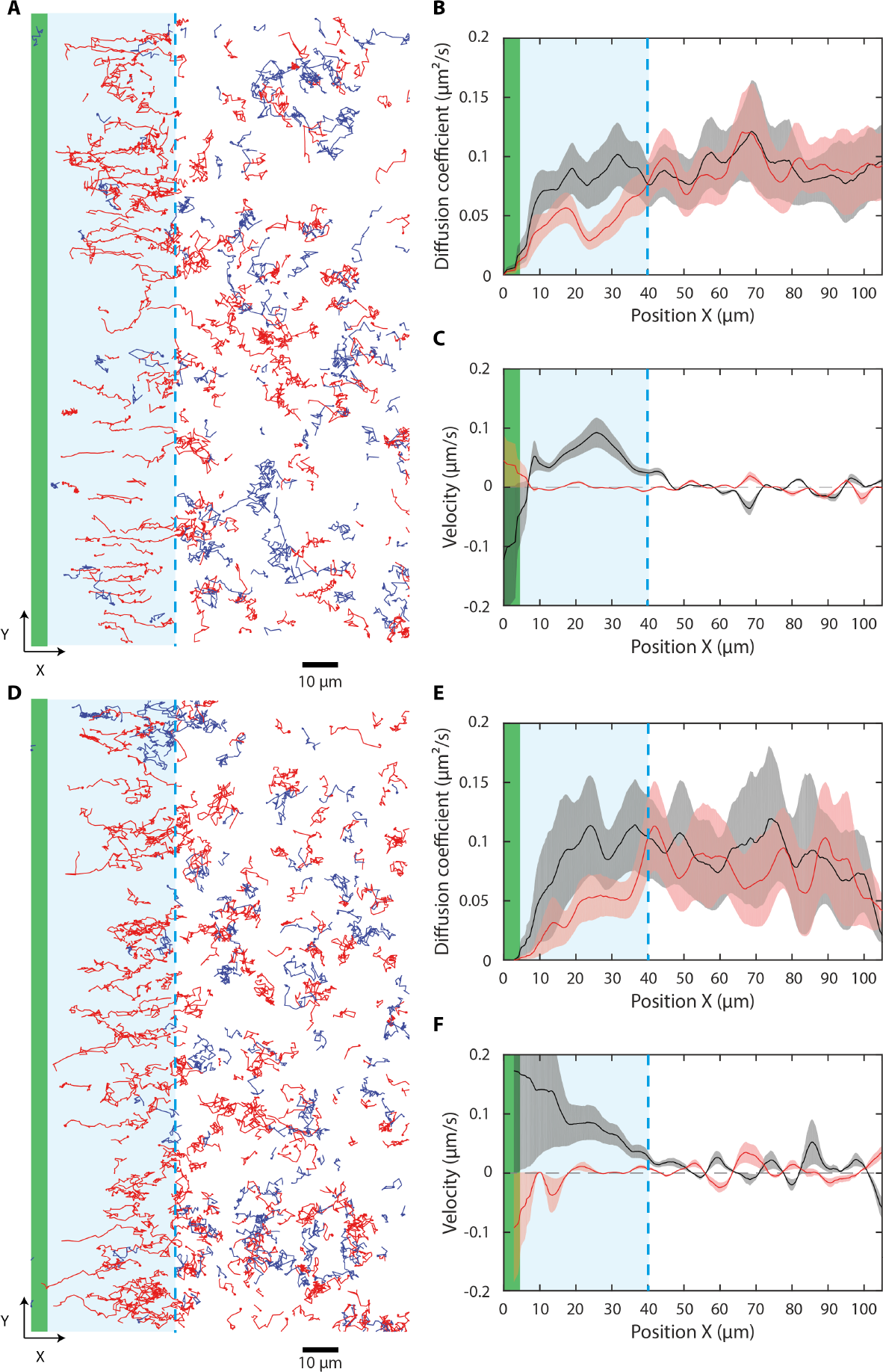
Bead transport in parallel actin networks. **A**: Trajectories of myosin-V-coated beads within the plane of the actin network. Directed movements oriented perpendicular to the nucleation line (green) and going away from the pattern are observed for distances *X* ≤ 40 *μ*m (blue-shaded area) from the nucleation line; at larger distances, the beads show diffusive movements. Red (blue): trajectories with a net positive (negative) movement along axis *X*. Diffusion coefficient (**B**), and velocity (**C**) of the beads as a function of bead position along axis *X* (black) and axis *Y* (red) for the data shown in **A**. **D-F**: Same as in **A-C** but for myosin-II-coated beads.

**FIG. S2:**
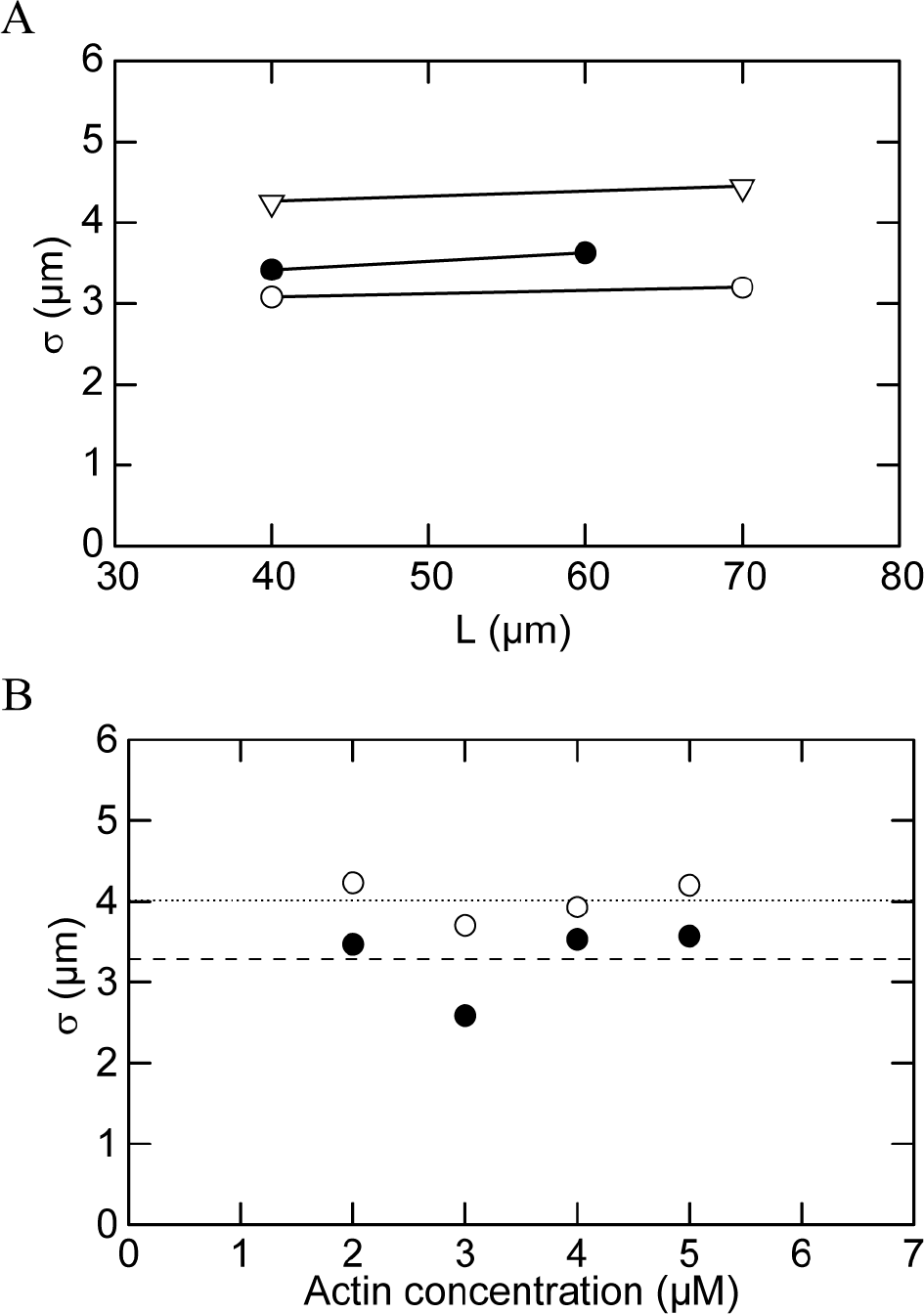
Effect of the actin-network size and of the actin concentration on bead positioning. The standard deviation *σ* at steady state is plotted as a function of the spacing *L* between nucleation lines of the actin micropattern for myosin-II-coated beads (**A**) and as a function of monomeric-actin concentration in the polymerization mix (**B**). The different symbols mark three different experiments in **A** and myosin-II-coated beads (white disks) or myosin-V-coated beads (black disks) in **B**.

### Section 2: Bead dynamics in antiparallel actin networks

We present here the physical model used to interpret the behavior of beads coated with either heavymero myosin-II or myosin-V motors moving on heterogeneous one-dimensional actin networks. Our description extends a theory developed originally to capture endosomal transport during the asymmetric division of sensory organ precursors in Drosophila [1]; we here consider slowly varying density profiles of antiparallel filament networks instead of overlapping homogeneous networks of opposite polarities. Single beads can be in one of the following three states: bound to actin filaments whose barbed end points rightwards (+) or bound to actin filaments whose barbed end points leftwards (−) or unbound (see main text, Fig. 5A). In the two bound states, beads move towards the barbed end of the filaments at constant speed *v* in the direction set by the polarity of the actin filament. In the unbound state, beads diffuse passively with a diffusion coefficient *D*_∅_. Transitions between states are characterised by three independent rate constants. A bead can switch from a bound state to the unbound state with a detachment rate constant *k*_OFF_ that does not depend on position or filament polarity. Alternatively, a bead can switch from the unbound state to a bound state with an attachment rate constant *k*_ON*ρ*_^+^ or *k*_ON*ρ*_^−^ that depends both _ON_ position and _ON_ the polarity of the filaments to which the bead binds. We define the local density *ρ*+ of filaments of polarity + (rightwards) and *ρ*− of filaments of polarity − (leftwards). To account for experimental observations (see main text, Fig. 1), we assume that the density profiles take the form *ρ*^+^ = (*ρ*_0_/2)*e*^−*X*/**λ**^ and *ρ*^−^ = (*ρ*_0_/2)*e*^*X*/**λ**^, where **λ** stands for the mean actin length and *ρ*_0_ = *ρ*^+^(*X* = 0) + *ρ*^−^(*X* = 0) is the total density of actin filaments at the center of the network (i.e. *X* = 0, where *ρ*^+^ = *ρ*^−^). Finally, the local probability distribution of beads bound on actin filaments oriented rightwards (leftwards) is denoted by *P*^+^ (*P*^−^) and the local probability distribution of unbound beads is denoted by *Pd*. The corresponding master equations that govern the stochastic dynamics of our beads read

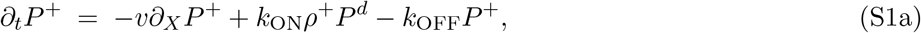

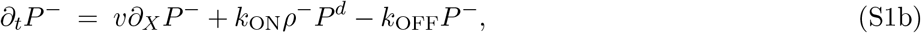

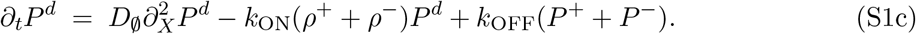

We performed simulations using a numerical scheme based on Gillespie’s First Reaction algorithm, [2]. To this end, we transform the spatial coordinate *X* into a discrete variable by introducing a step size Δ*X*. In the bound state, beads advance a unit step Δ*X* towards the barbed end of the actin filament at a rate constant *v*/Δ*X*. As a result of velocity fluctuations in the bound states, discretization introduces an effective diffusion coefficient that scales as ~ *v*Δ*X*. The value of Δ*X* used in the simulations (0.05 *μ*m) ensured that this contribution to the diffusive motion of the beads produced a small deviation (< 10%) to the actual bead diffusion coefficient. The diffusive motion of the beads in the unbound state is described as a simple random walk with a hoping rate constant *D*_∅_/Δ*X*^2^. Similarly, *k*_OFF_ and *k*_ON_*ρ*^±^ control the transitions between the three states of a bead, as explained above. We choose reflective boundary conditions, enforcing that the total number of particles remains constant over time.

In this model, we can define two intrinsic length scales: the *ballistic length lB* = *v/k*_OFF_ and the *diffusive length* 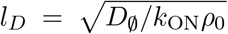, which control the mean distance traveled during runs in a bound state and the mean-squared displacement of diffusive motion in the unbound state, respectively. Additionally, we can define two time scales: the attachment time *τ*_ON_ = 1/*k*_OFF_ and the detachment time *τ*_OFF_ = 1/(*k*_ON_*ρ*_0_), which control the mean time spend by the bead in the bound and in the unbound states, respectively. Consequently, model-bead behaviors can be classified in four asymptotic regimes, depending on the dominant characteristic time and length scale in the system.

**FIG. S3:**
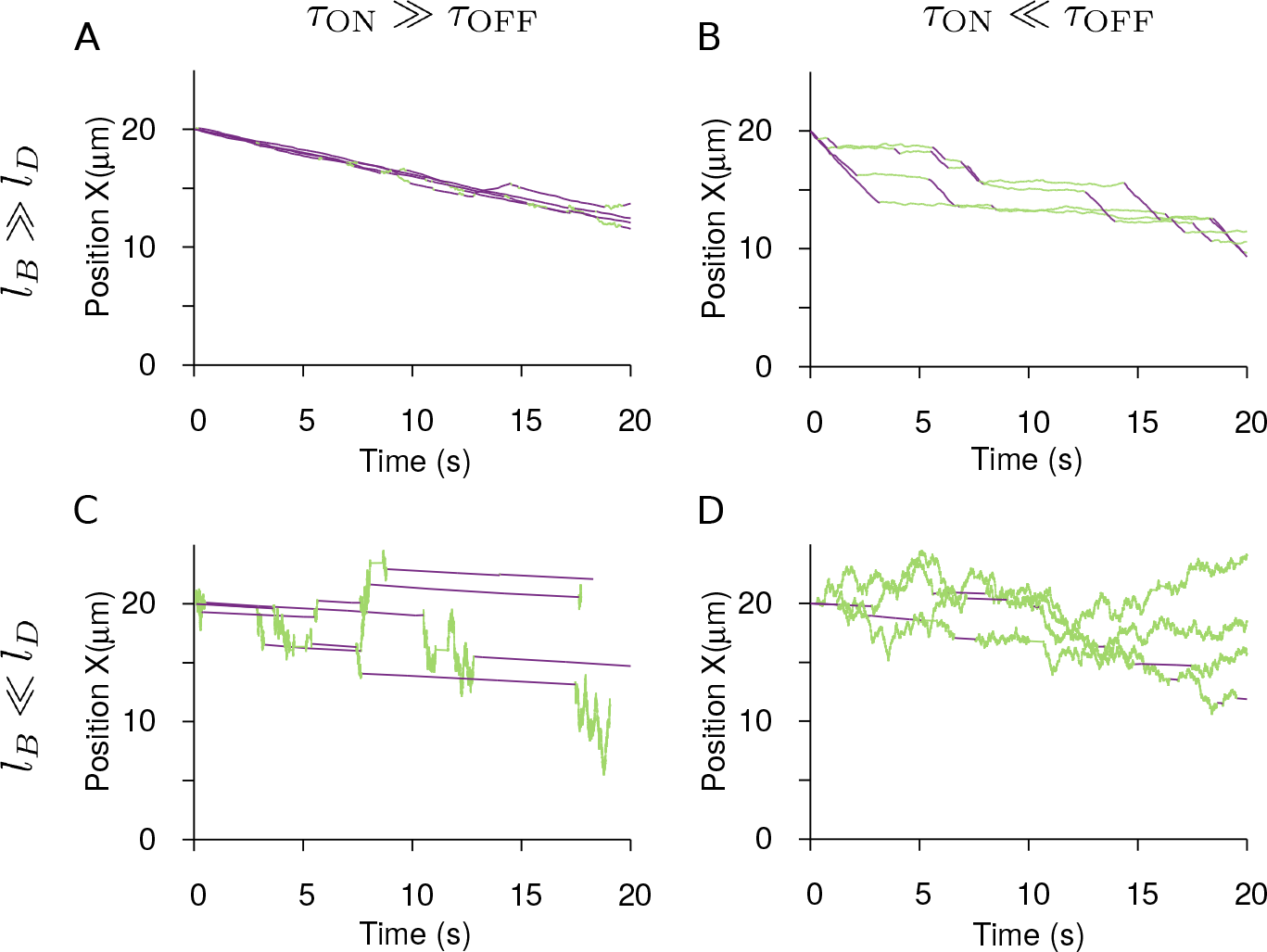
Short-time dynamics in the four limiting regimes of bead transport. In (**A** and **B**) *l*_*B*_ ≫ *l*_*D*_, whereas in (**C** and **D**) *l*_*B*_ ≪ *l*_*D*_. In (**A** and **C**) *τ*_ON_ ≫ *τ*_OFF_, whereas in (**B** and **D**) *τ*_ON_ ≪ *τ*_OFF_. The magenta (green) segments of the trajectories mark when the bead is bound (unbound). The list of parameters is given in Table S1.

In Figure S3, we show simulations of the short-time dynamics of model beads initially located at a 20-*μ*m distance (*X* = 20 *μ*m) from the geometrical center (*X* = 0) for all four asymptotic regimes. When *l*_*B*_ ≫ *l*_*D*_, bead transport is dominated by the ballistic motion in the bound states. Qualitatively, the trajectories are composed of an alternation of pauses in the unbound state and runs in the bound states (Fig. S3A-B). When *l*_*B*_ ≪ *l*_*D*_, the passive diffusive motion in the unbound state instead dominates the motion of the beads. Qualitatively, in this case the trajectories are composed of an alternation of diffusive motion in the unbound state and pauses in the bound states (Fig. S3C-D). When *τ*_ON_ ≫ *τ*_OFF_, the beads are mostly bound (Fig. S3A,C) and when *τ*_ON_ ≪ *τ*_OFF_, the beads are mostly unbound (Fig. S3B,D).

**Fig. S4.**
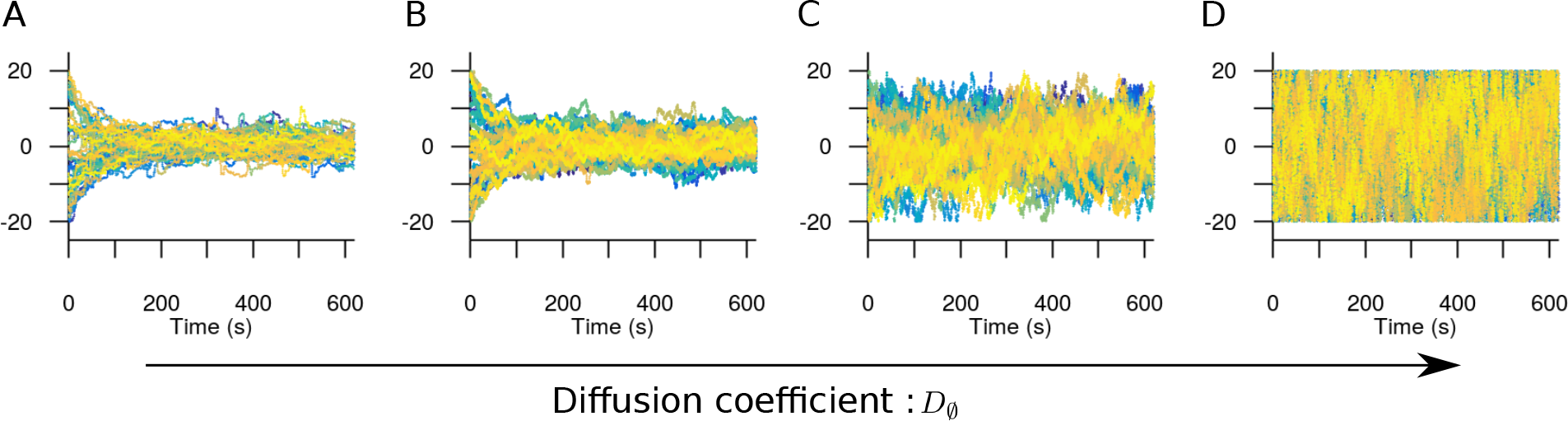
Long-time bead dynamics. The diffusion coefficient *D*_∅_ is: (**A**) 7 ∗ 10^−3^, (**B**) 7 ∗ 10^−2^, (**C**) 7 ∗ 10^−1^ and (**D**) 7, in unit of *μ*m^2^/*s*. The diffusive length *l*_*D*_ is: (**A**) 0.2, (**B**) 0.7, (**C**) 2 and (**D**) 7, in unit of *μ*m, whereas the ballistic length *l_B_* = 0.7 *μ*m is fixed. Other parameters are listed in Table S2 (Case 1).

In Figure S4, we show the Long-time dynamics of the beads as a function of the diffusion coefficient *D*_∅_; the beads are initially in a random state and distributed randomly in the network *X* ∈ (+20, −20) *μ*m. We find that the beads move, on average, towards the center of the network, where the densities of both types of actin filaments are equal and the net polarity of the network thus vanishes. As one would expect, increasing the diffusion coefficient *D*_∅_ results in larger fluctuations in the unbound state and a broadening of the range of positions explored by the beads at steady state. Remarkably, the amplitude of these fluctuations does not vanish when the diffusion coefficient goes to zero. As will be clarified below, these fluctuations are the consequence of the active stochastic dynamics of the beads when they are bound to the filaments, which generates noise.

### Section 3: Effective velocity and diffusion coefficient

In this Section, we discuss simulations of the Long-time behavior of the mean and the mean-squared displacement of bead assemblies. We show that a local velocity and diffusion coefficient, characteristic of bead trajectories in heterogeneous networks, can be defined unequivocally when the scale of the heterogeneities in the actin network is large *λ* ≫ *l*_*B*_, *l*_*D*_. Based on the profiles of bead velocity and diffusion coefficient profiles that were measured in antiparallel actin networks, we conclude that myosin-coated bead transport is constrained to the regime where the beads detach often *τ*_ON_ ≪ *τ*_OFF_ and travel comparable distances in the attached and detached states *l*_*D*_ ~ *l*_*B*_.

When *λ* ≫ *l*_*B*_, *l*_*D*_, we can assume local uniformity of the actin network, measuring that the actin density profiles *ρ*^+^ and *ρ*^−^ are uniform or space-independent: the beads experience a “uniform” actin density during the relaxation of their internal degrees of freedom, which occurs over a time scale set by *τ*_ON_ and *τ*_OFF_.

Let’s start with all beads initially in the unbound state (*P* (*t* = 0) = *P*^*d*^(*t* = 0) = *δ*(*X* − *X*_0_)), where *X*_0_ is the initial location and *P* = *P*^+^ + *P*^−^ + *P*^*d*^ is the total probability distribution of beads (see Eq. S1). Without loss of generality, we take *X*_0_ = 0 since the actin network is uniform. The mean displacement and the mean squared displacement are defined as 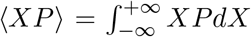 and 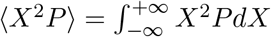, where the brackets refer to the integration over the whole spatial domain.

We perform the Laplace transformation in time on Equations S1 with 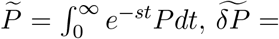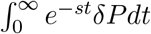 and 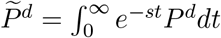, leading to

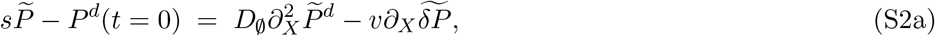

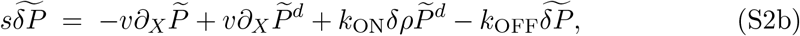

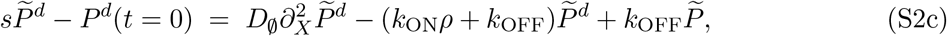

where *ρ* = *ρ*_+_ + *ρ*_−_ is the total density of actin filaments, *δ*_*ρ*_ = *ρ*_+_ − *ρ*_−_ and *δP* = *P*^+^ − *P*^−^. From Equations S2, we deduce that the distribution of beads, the mean displacement and the mean squared displacement in Laplace space obey

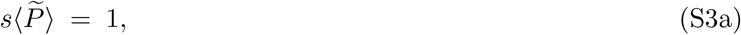

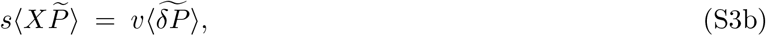

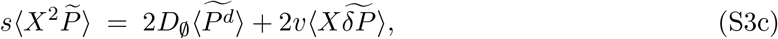

from which we obtain analytically their time evolution. The boundary terms of the form ~ *X*^*n*^*P*|_±∞_ have been set to zero regardless of the probability distribution.

After integrating Equations S2 over space, we obtain the following equations

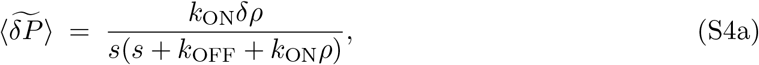

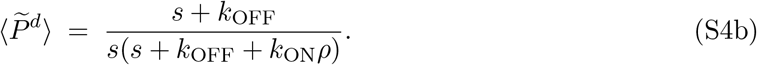

Similarly, multiplying first by *X* and then integrating Equations S2 over space, the following equations are obtained

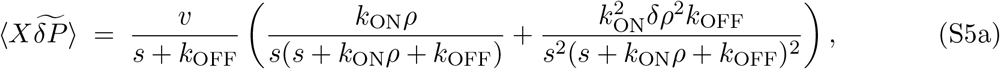

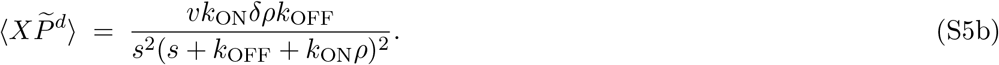

Inserting them back into Equations S3b-S3c, one deduces an explicit form of the mean displacement and mean square displacement that read

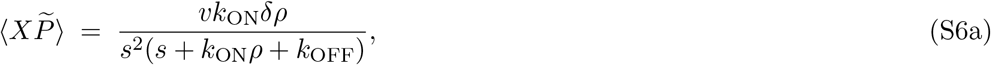

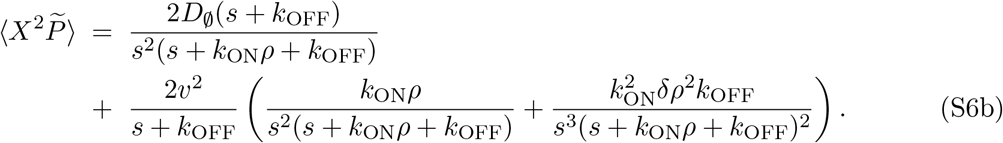

By applying the inverse Laplace transformation to Equations S6, we derive the time evolution of these observables. Here, we focus on the Long-time behavior of the mean displacement (S6a) and of the mean squared displacement (S6b), which is achieved through the limit *t* → ∞. In the Long-time limit, we define an effective local velocity and diffusion coefficient as the coefficient proportional to time of the mean displacement and variance, respectively. From Equations S6, we obtain

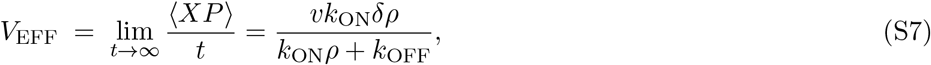

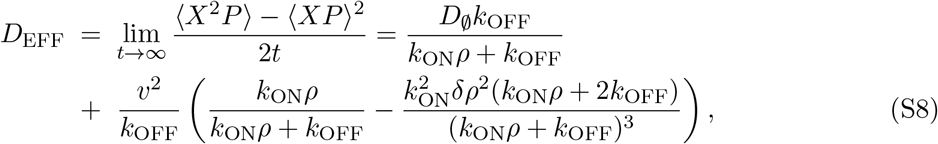

which are valid at any location in the actin network (i.e. for any functional form of *ρ*^+^(*X*) and *ρ*^−^(*X*)) and at time scales Longer than the relaxation time scale of the internal degrees of freedom of the beads (i.e. *t* ≫ *τ*_ON_ and *t* ≫ *τ*_OFF_). The validity of the previous equations requires local uniformity of the actin network, meaning that *λ* ≫ *V*_*EFF*_/*k*_OFF_ and **λ**^2^ ≫ *D*_*EFF*_/*k*_OFF_ or equivalently *λ* ≫ *l*_*B*_, *l*_*D*_. These Equations S7-S8 have also been derived from the relaxation of the hydrodynamical modes in Ref. [1].

Note that Equations S7-S8 are in general controlled by both the physical properties of molecular motors and by the architecture of the underlying actin network. Interestingly, the effective velocity (S7) is proportional to the difference in the densities of filaments of opposite polarities *δ*_*ρ*_ = *ρ*^+^ −*ρ*^−^, meaning that beads are directed towards regions where *δ*_*ρ*_ = 0 or equivalently where there is no net polarity. With respect to the diffusion coefficient, Equation S8 reflects the presence of two fundamentally different mechanisms of diffusive-like motion. The first term of Equation S8, proportional to *D*_∅_, arises from the passive diffusion of beads in the unbound state. Instead, the second term of Equation S8, proportional to *v*^2^/*k*_OFF_, arises from the persistent random walk dynamics performed by beads in the bound states. Specifically, near the center, where *δ*_*ρ*_ = 0, the step size of this random walk is the run length *l*_*B*_ = *v*/*k*_OFF_ and the switching rate constant is *k*_OFF_, so that the effective diffusion of our random walkers would be 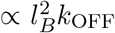 times the steady state probability of beads in the bound state (*k*_ON*ρ*_/(*k*_ON*ρ*_ + *k*_OFF_)). The beads thus undergo both passive and active diffusion.

In Figure S5, we show the profiles of the effective velocity *V*_EFF_(*X*) and effective diffusion coefficient *D*_EFF_(*X*) (S7-S8), assuming that *ρ*_+_ = (*ρ*_0_/2)*e*^−*X*/**λ**^ and *ρ*_−_ = (*ρ*_0_/2)*e*^*X*/**λ**^, as in experiments (see Fig. 1 of the main text). The competition between the two time scales strongly affects bead motion (Fig. S3) and their effects are reflected in *V*_EFF_(*X*) and *D*_EFF_(*X*) as well (Fig. S5). In the limit of low attachment rates (*τ*_ON_ ≪ *τ*_OFF_), the effective velocity and effective diffusion coefficient (S7-S8) can be approximated by

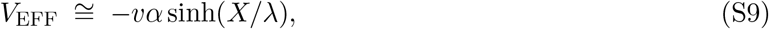

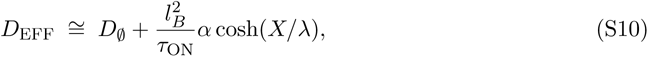

where *α* = *τ*_ON_*/τ*_OFF_. In the opposite limit, high attachment rates (*τ*_ON_ ≫ *τ*_OFF_), the same effective velocity and effective diffusion coefficient can be approximated by

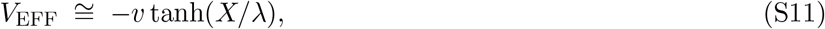

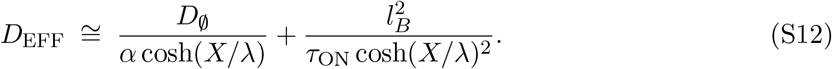

In the limit *τ*_ON_ ≫ *τ*_OFF_, the effective velocity near the center is well approximated by a linear function of *X* with a slope equal to *v*/**λ** (Fig. S5A), whereas in the limit *τ*_ON_ ≪ *τ*_OFF_, the slope drops to *vτ*_ON_/(*τ*_OFF_**λ**) < *v*/**λ** (Fig. S5B). Experimentally, we observed that myosin-II-coated and myosin-V-coated beads exhibit mean velocities that are on average smaller than the corresponding speed *v* of molecular motors in motility assays [3]. By fitting this expression to the actual slope of the velocity profiles (see Fig. 4 of the main text), we find that for myosin-V-coated beads *τ*_ON_*/τ*_OFF_ = 0.05 with *v* = 2 *μ*m/s, and for myosin-II-coated beads *τ*_ON_*/τ*_OFF_ = 0.17 with *v* = 0.46 *μ*m/s. Thus, suggesting that near the center, beads operate in the regime *τ*_ON_ ≪ *τ*_OFF_, or equivalently beads detach often.

**Fig. S5.**
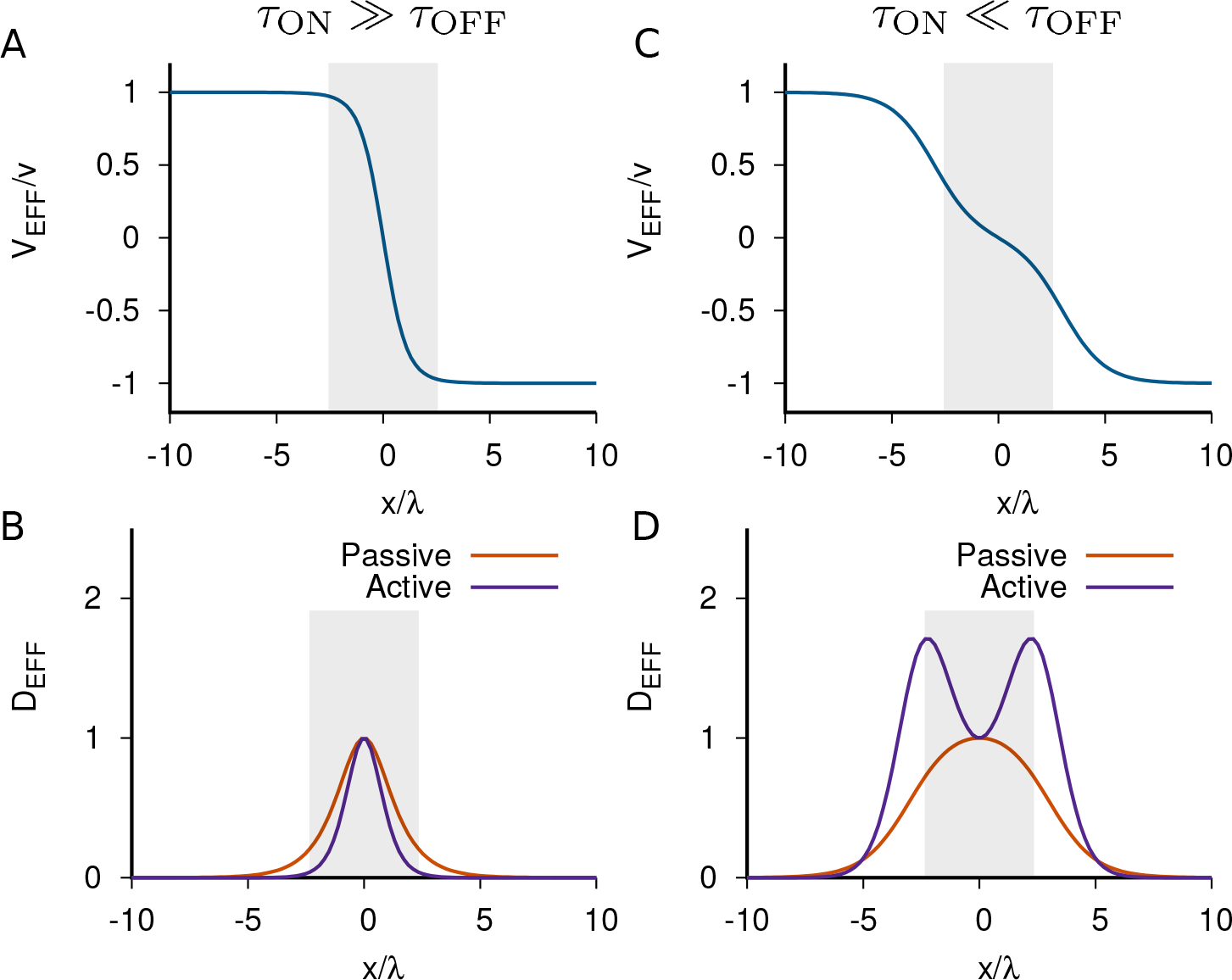
Effective velocity and effective diffusion coefficient profiles (S7-S8) in the two limiting regimes set by the competition between the time scales *τ*_ON_ and *τ*_OFF_. (**A** and **B**) represent the normalized velocity and diffusion profiles, respectively, for a value of *τ*_ON_*/τ*_OFF_ = 10. (**C** and **D**) represent the same profiles but for a value of *τ*_ON_*/τ*_OFF_ = 0.1. The velocity (blue) has been normalized by *v* and each contribution to the diffusion coefficient has been normalized by their value at the center. The passive (active) contribution of the diffusion coefficient is shown in red (purple). The gray shaded area represents the region in between two nucleation lines *L* = 40 *μ*m.

The effective diffusion coefficient (S8) originates from two independent mechanisms: passive and active diffusion. The passive mechanism (red curves) produces an effective diffusion constant with a peak at the center, regardless of the dominant time scale (Figure S5C-D). Instead, the active diffusion (purple curves) yields a profile of the diffusion coefficient with a shape that depends on the dominant time scale: when *τ*_ON_ ≫ *τ*_OFF_, there is a maximum at the center (Fig. S5C), whereas when *τ*_ON_ ≪ *τ*_OFF_, there is a minimum at the center, akin to passive diffusion (Fig. S5D). Experimentally, the measured diffusion coefficient profiles of myosin -II and -V beads display a marked minimum near the center (see Fig. 4 of the main text), suggesting that diffusion is dominated by the contribution resulting from active bead transport. Specifically, the curvature of the effective diffusion coefficient profile (S8) is positive when 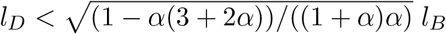 in which *α* = *τ*_ON_*/τ*_OFF_. For myosin-V-coated beads, with *α* = 0.05, this condition reads *l*_*D*_ < 4*l*_*B*_ and for myosin-II-coated beads, with *α* = 0.17, *l*_*D*_ < 1.5*l*_*B*_. Because the ratio of passive and active contributions to the effective diffusion coefficient (Eq. S8) is given by (*l*_*D*_/*l*_*B*_)^2^, these constraints enforce that at the center of anti-parallel networks, the passive contribution is at most 16 (2) times larger than the active contribution for myosin-V-coated (myosin-II-coated) beads (Eq. S8). To determine the diffusion coefficient *D*_∅_, and by extension the time scales *τ*_ON_ and *τ*_OFF_, we need to introduce an additional constraint. By assuming that *D*_∅_ = 0 and using the values of Fig. 4H in the main text to determine the effective diffusion coefficient (S8) at the center, we obtain the set of parameters listed in Table 1 of the main text. Of note, the absolute values of the time scales *τ*_ON_ and *τ*_OFF_ are model dependent. If we instead assume that the two length scales are equal *l*_*B*_ = *l*_*D*_ (i.e. *D*_∅_ = *v*^2^*α*/*k*_OFF_), a condition that leads to nearly optimal centering (see Fig. 5 of the main text), then the value of the time scales are halved with respect to the parameters in Table 1 (Table S2). In either case, we can conclude that myosin-coated bead transport operates in a regime where the beads detach often (*τ*_ON_ ≪ *τ*_OFF_).

At this point, we wondered whether the steady-state statistics of beads in anti-parallel networks can be described by an advection-diffusion process, as in experiments (see Fig. 4C,F in the main text). To address this question, we computed the steady-state bead distribution by solving numerically the stochastic dynamics of ~ 100, 000 beads in antiparallel networks for each condition (black curve in Fig. S6) and compared this distribution to that resulting from a diffusion-drift process *P*_*DD*_ ∝ exp(*∫*_*X*_ *V*_EFF_(*y*)/*D*_EFF_(*y*)*dy*) (green curve in Fig. S6) with the velocity and diffusion coefficients (S7,S8). Fig. S6 shows an excellent agreement between both distributions for a broad range of parameters extending both towards the large diffusion limit *l*_*D*_ ≫ *l*_*B*_ and the low diffusion limit *l*_*D*_ ≪ *l*_*B*_. In conclusion, at steady state, the model reproduces robustly the advection-diffusion approximation of the statistics of motor-coated beads.

The distribution of model beads peaked in the center (*X* = 0, where *ρ*^+^ = *ρ*^−^): the active transport process brought the beads, on average, to the position where the net polarity of the network vanishes. Near the center, Equations (S7,S8) admits a Taylor expansion

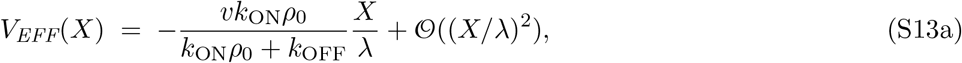

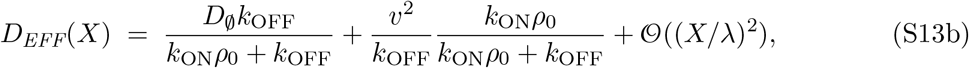

which is valid only if the mean length of actin filaments *λ* is much larger than *X*. Interestingly, these expressions lead to an approximate Gaussian distribution of model-bead position with a peak located at the geometrical center *X* = 0 and a variance

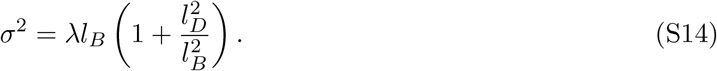

**Fig. S6.**
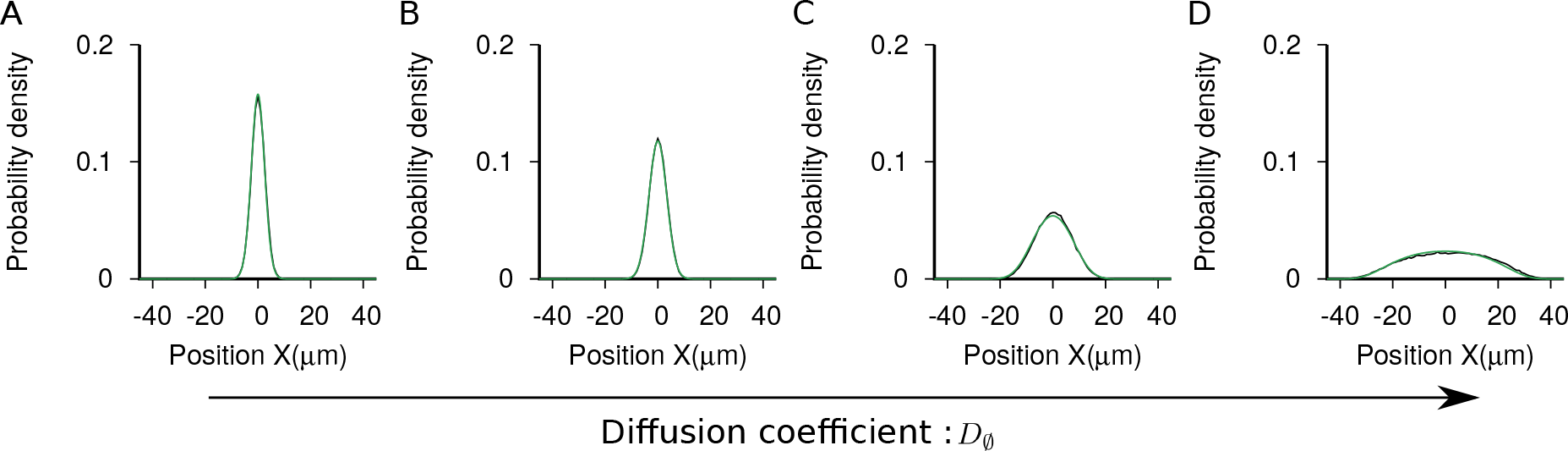
Validity of the approximation of model-bead statistics by a diffusion-drift process. The black curve corresponds to the steady-state bead distribution *P* (*X*) resulting from solving Equations S1 in an antiparallel network. The green curve is the probability distribution expected from a diffusion-drift process *P*_*DD*_ ∝ exp(*∫*_*X*_ *V*_EFF_(*y*)/*D*_EFF_(*y*)*dy*) with the velocity and diffusion coefficient (S7,S8). The diffusion coefficient *D*_∅_ is: (**A**) 7 ∗ 10^−3^, (**B**) 7 ∗ 10^−2^, (**C**) 7 ∗ 10^−1^ and (**D**) 7, in unit of *μ*m^2^/*s*. The diffusive length *l*_*D*_ is: (**A**) 0.2, (**B**) 0.7, (**C**) 2 and (**D**) 7, in unit of *μ*m, whereas the ballistic length *l*_*B*_ = 0.7 *μ*m is fixed. Other parameters are listed in Table S2 (Case 1).

Thus, our model of active bead transport describes positioning at the center of the network with a precision approximated by Equation S14 up to a numerical pre-factor. We note that the theory predicts that the variance of the bead position at steady state does not depend on the size *L* of the network, meaning that bead centering depends on the local architecture of the network in a neighborhood centered where the net polarity vanishes. Rather, the precision of positioning is set by the ballistic length *l*_*B*_, the diffusive length *l*_*D*_, and the length scale *λ* of heterogeneities within the network. Remarkably, by systematically increasing *l*_*B*_ in Equation S14, we find that the variance (S14) of model-bead position at steady state first decreases but then reaches a minimum at *l*_*D*_ = *l*_*B*_, where the variance is 2**λ*l*_*B*_, corresponding to an optimum of centering precision (Fig. 5H-I in the main text). When *l*_*B*_ ≪ *l*_*D*_, bead displacements are largely due to the diffusive motion in the unbound state and the variance near the center scales as 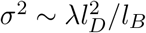. As expected increasing the diffusive length *l*_*D*_ leads to poor centering. When *l*_*B*_ ≫ *l*_*D*_, bead displacements are largely due to the ballistic runs in the attached states and the variance near the center scales as *σ*^2^ ~ **λ*l*_*B*_. In this limit, centering is only controlled by two of the three characteristic lengths (**λ**, *l*_*B*_). Decreasing the ballistic length *l*_*B*_ leads to better centering, suggesting that in the absence of passive diffusion (*D*_∅_ = 0), perfect centering (i.e. *σ* = 0) is achieved only when the ballistic length *l_B_* = 0 or in other words when motor speed vanishes *v* = 0. These results are in qualitative agreement with numerical estimates (Fig. 5H-I in the main text). The numerical relation between the variance model-bead position at steady state and the ballistic length *l*_*B*_ shows a minimum; the minimum occurs here at *l*_*D*_ ≈ 1.65*l*_*B*_, where the variance is 2.07**λ*l*_*B*_. In conclusion, model beads are able to position themselves where the net polarity of the network vanishes with a precision that exhibits an optimal value when the two characteristic length scales of bead dynamics are comparable (*l*_*B*_ ~ *l*_*D*_).

### Section 4: Effects of filament switching on bead dynamics

In the preceding Sections, we assumed that a bead could be bound to one type of filament (+ or −) only. Attachment to filaments of opposite polarity can result in a tug-of-war. Because tug-of-war is mechanically unstable [4], a bead engaged in this situation is expected to switch direction by changing the filaments to which it is bound while remaining bound. We describe this process by a direct stochastic transition of beads between the two bound states without passing through an intermediate unbound state. We assume that beads switch between the bound states at a rate constant *kρ*^±^ that depends proportionally on the actin density of the filament type to which beads bind. The corresponding master equations that govern the stochastic dynamics of our model beads now read

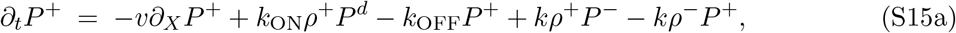

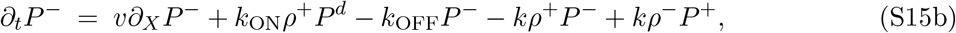

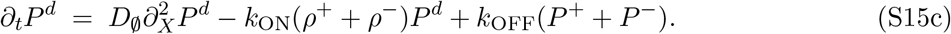

Above, we have discussed a mechanism harbored in Equations S1 (i.e. Eq. S15 when *k* = 0) by which beads reach the center (where *ρ*^+^ = *ρ*^−^) of anti-parallel networks. Beads are able to sample asymmetries in actin densities by combining directed motion when bound plus detachment-attachment cycles. We name this mechanism *detachment-based centering*. Equations S1, however, omit the direct transitions between actin filaments of different type. Here, we explore how filament switching affects the bead dynamics.

First, we wonder about the effects of filament switching for bead transport in anti-parallel actin networks, when detachments are allowed *k*_OFF_ ≠ 0. In Figure S7, we show the influence of switching for both the Long-time behavior of bead trajectories and the steady-state distribution of beads. We observe that increasing the switching rate constant, leads to enhanced funneling of the trajectories (Fig. S7A-C) and narrowing of the bead distribution (Fig. S7D-F). Thus, switching favors centering of beads by sharpening detachment-based centering.

**Fig. S7.**
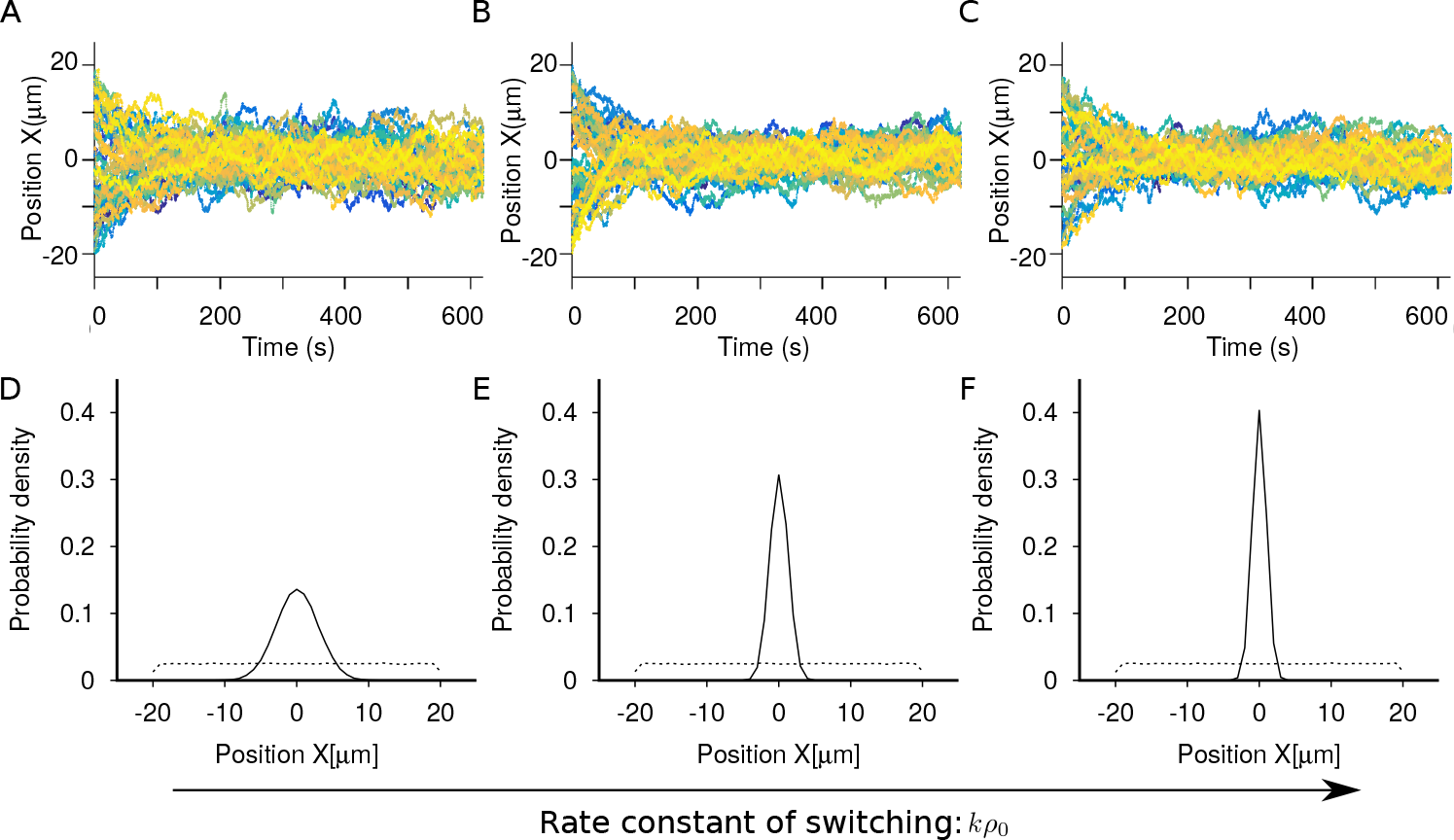
Effects of switching on bead transport. (**A-C**) show the Long time dynamics of beads, and (**D-F**) show the total distribution of beads at steady state (solid black) and at the initial instant (dashed black). The parameters are listed in Table S2 (Cases 1). The switching rate constant increases: *kρ*_0_ = 0 (**A** and **D**), *kρ*_0_ = 6 (**B** and **E**) and *kρ*_0_ = 12 (**C** and **F**).

Generalizing the derivation shown in Section 3 to systems with switching results in bead distributions that peak at the center of the network with a variance 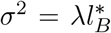 that depend on the renormalized ballistic length 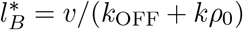. This result is in agreement with Figure S7, for the increase of the switching rate constant leads to the sharpening of the bead distribution. Thus, switching effectively decreases the ballistic length, which now can be interpreted as the mean distance that a bead moves in the same direction when attached to actin, resulting in finer sampling of the net-polarity gradient and more precise positioning at the center.

Second, we wondered whether switching alone (i.e. in the absence of detachment events *k*_OFF_ = 0) is sufficient to center beads in antiparallel actin networks, where the actin densities takes the usual form *ρ*^+^ = (*ρ*_0_/2)*e*^−*X*/**λ**^ and *ρ*_ − = (*ρ*_0_/2)*e*^*X*/**λ**^. In Figure S8, we present the statistical properties of bead trajectories, when detachment is absent (Fig. S8A-C) or when switching is absent (Fig. S8D-F). The latter corresponds to the situation studied above. Remarkably, we find that similar (i.e. peaked) steady-state distribution of model beads can be achieved when there is only switching (Fig. S8C,F), meaning that this mechanism is sufficient to induce bead centering. The typical trajectories of switching-based transport are composed of an alternation of runs between the two types of filaments (Fig. S8A,B). Note that the trajectories of detachment-based transport are composed of a combination of pauses in the unbound state and runs in the bound state (Fig. S8D,E).

**Fig. S8.**
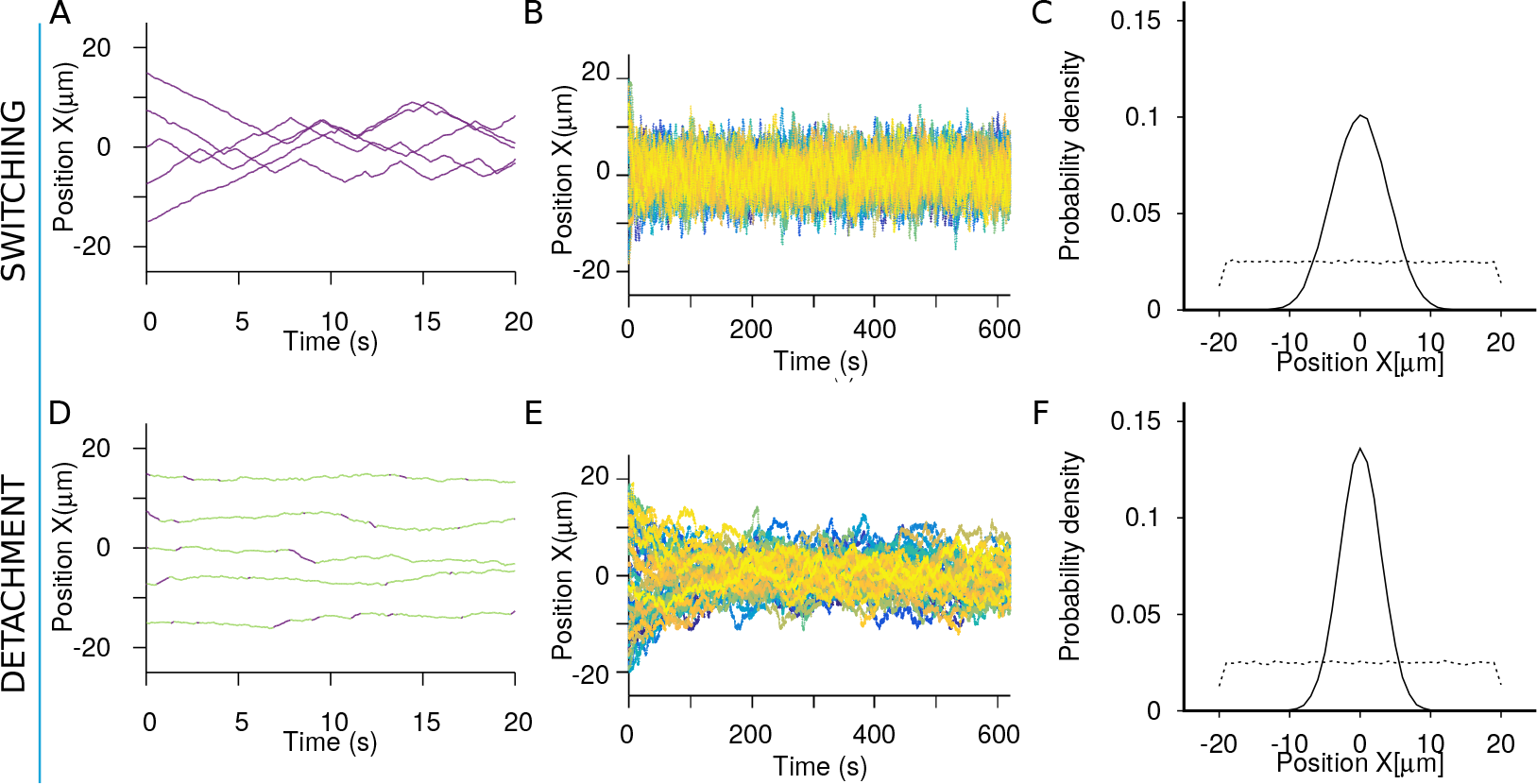
Comparison between detachment-based and switching-based bead transport. (**A** and **D**) and (**B** and **E**) show the short and Long time dynamics of beads, respectively. (**C** and **F**) show the total distribution of beads at steady state (solid black) and at the initial instant (dashed black). The parameters are listed in Table S2 (Cases 1). In panels (**A-C**) the detachment rate constant *k*_OFF_ = 0 and the switching rate constant *kρ*_0_ = 0.5, and in panels (**D-F**) *k*_OFF_ = 2.9 and *kρ*_0_ = 0.

Without going into details, similar calculations as those described in Section 2 lead to an effective velocity and diffusion coefficient for the transport of beads, when detachment is absent *k*_OFF_ = 0

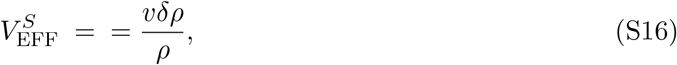

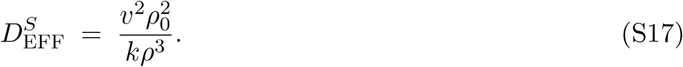

Velocity and diffusion profiles given by Eqs. S16-S17 are shown in Fig. S9 for anti-parallel actin networks with the center at *X* = 0. Note, that neither the slope of the velocity at the center (~ *v*/**λ**) nor the convex shape of the diffusion coefficient agrees with our experimental observations on myosin-II and -V-coated beads, suggesting that switching is not the mechanism at work.

**Fig. S9.**
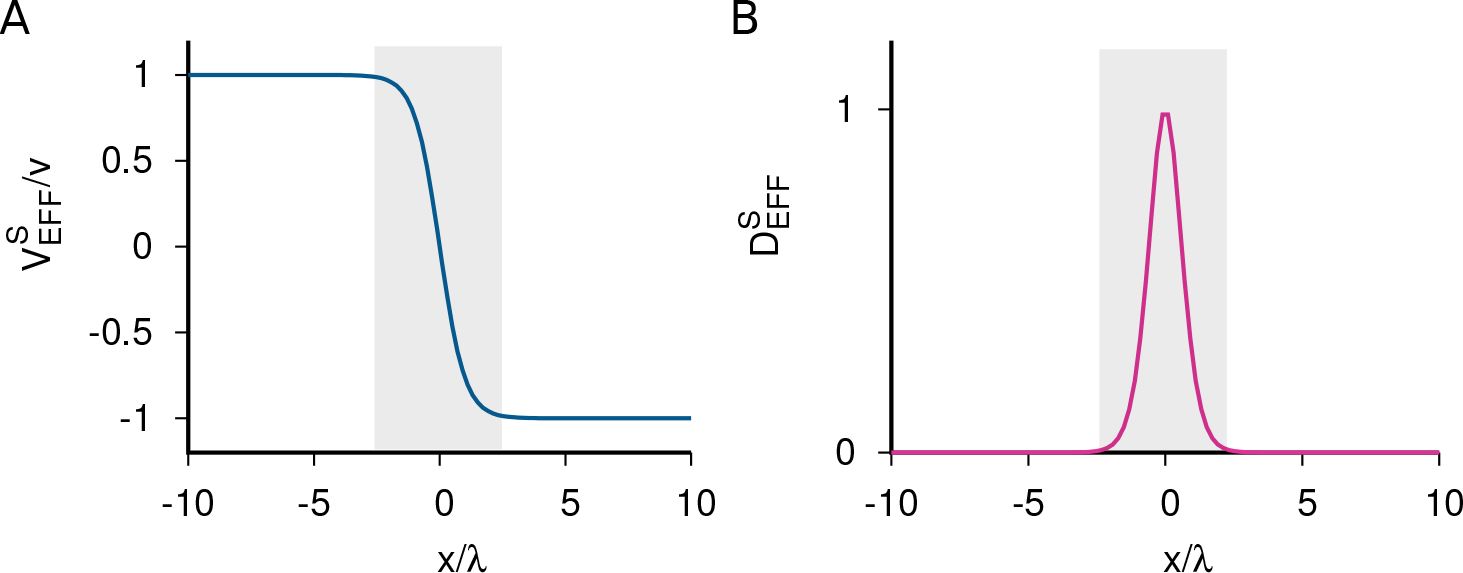
Representation of the effective velocity (**A**) and diffusion coefficient (**B**) given by equations (S16-S17) as a function of the normalized position with respect to the mean actin length **λ**. The velocity (blue) has been normalized by *v* and the diffusion coefficient (pink) has been normalized by *v*^2^/*kρ*_0_. The gray shaded area represents the space in between two nucleation lines *L* = 40 *μ*m.

### Section 5: Tables of parameters

**TABLE S1:** List of parameters used to produce the stochastic trajectories shown in Fig. S3.

| | $k_{\text{ON}}\rho_0[1/s]$ | $k_{\text{OFF}}[1/s]$ | $v[\mu\text{m}/s]$ | $D_\emptyset[\mu\text{m}^2/s]$ | $L[\mu\text{m}]$ | $\lambda[\mu\text{m}]$ |
| --- | --- | --- | --- | --- | --- | --- |
| Fig. (1,a) | 0.8 | 0.4 | 0.4 | 0.1 | 40 | 8.7 |
| Fig. (1,b) | 0.08 | 1 | 2 | 0.01 | 40 | 8.7 |
| Fig. (1,c) | 0.8 | 0.4 | 0.1 | 10 | 40 | 8.7 |
| Fig. (1,d) | 0.08 | 1 | 0.1 | 1 | 40 | 8.7 |

**TABLE S2:**
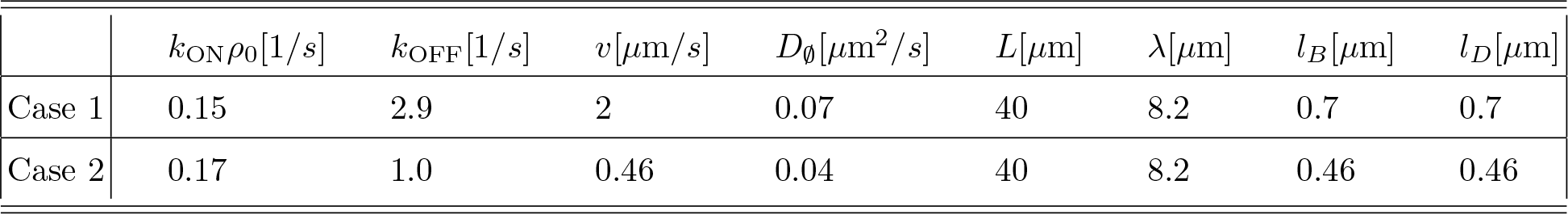
Parameters of simulations describing experimental bead trajectories.

| | $k_{\text{ON}}\rho_0[1/s]$ | $k_{\text{OFF}}[1/s]$ | $v[\mu\text{m}/s]$ | $D_\emptyset[\mu\text{m}^2/s]$ | $L[\mu\text{m}]$ | $\lambda[\mu\text{m}]$ | $l_B[\mu\text{m}]$ | $l_D[\mu\text{m}]$ |
| --- | --- | --- | --- | --- | --- | --- | --- | --- |
| Case 1 | 0.15 | 2.9 | 2 | 0.07 | 40 | 8.2 | 0.7 | 0.7 |
| Case 2 | 0.17 | 1.0 | 0.46 | 0.04 | 40 | 8.2 | 0.46 | 0.46 |

In Table S2, the speed *v* is set by the values obtained in motility assays [3]. The two length scales *l*_*B*_ and *l*_*D*_ are assumed to be equal. The distance between parallel nucleation stripes is an external controllable parameter with a typical value of 40 *μ*m. Case 1 (2) corresponds to the set of parameter estimated from the data in Fig. 4 in the main text for myosin-II-coated (myosin-V-coated) beads by using Equations (S7,S8).

